# GproDIA enables data-independent acquisition glycoproteomics with comprehensive statistical control

**DOI:** 10.1101/2021.03.20.436117

**Authors:** Yi Yang, Weiqian Cao, Guoquan Yan, Siyuan Kong, Mengxi Wu, Pengyuan Yang, Liang Qiao

**Affiliations:** Department of Chemistry, Institutes of Biomedical Sciences, and Shanghai Stomatological Hospital, Fudan University, Shanghai 200000, China; The Fifth People’s Hospital of Fudan University and the Shanghai Key Laboratory of Medical Epigenetics and the International Co-laboratory of Medical Epigenetics and Metabolism, Ministry of Science and Technology, Fudan University, Shanghai, China; NHC Key Laboratory of Glycoconjugates Research, Fudan University, Shanghai, China

## Abstract

Large-scale profiling of intact glycopeptides is critical but challenging in glycoproteomics. Data independent acquisition (DIA) is an emerging technology with deep proteome coverage and accurate quantitative capability in proteomics studies, but is still in the early stage of development in the field of glycoproteomics. We propose GproDIA, a framework for the proteome-wide characterization of intact glycopeptides from DIA data with comprehensive statistical control by a 2-dimentional false discovery rate approach and a glycoform inference algorithm, enabling accurate identification of intact glycopeptides using wide isolation windows. We further adapt a semi-empirical spectrum prediction strategy to expand the coverage of spectral libraries of glycopeptides. We benchmark our method for N-glycopeptide profiling on DIA data of yeast and human serum samples, demonstrating that DIA with GproDIA outperforms the data dependent acquisition (DDA) based methods for glycoproteomics in terms of capacity and data completeness of identification, as well as accuracy and precision of quantification. We expect that this work can provide a powerful tool for glycoproteomic studies.

## Main

Protein glycosylation is one of the most abundant and heterogeneous post-translational modifications (PTMs) that provides great proteomic diversity and plays a key role in various biological processes^1–3^, even the host–pathogen interaction of the ongoing coronavirus disease 2019 pandemic^4^. Precise characterization of protein glycosylation is critical for understanding mechanism of diseases^5,6^, discovery of biomarkers for diagnosis^7^, and development of drugs and vaccines^8^. The high heterogeneity of glycans across glycosites results in an increased number of glycoproteoforms. Profiling of intact glycopeptide provides the opportunity of simultaneous analysis of glycans, glycosite occupancy and site-specific glycosylation on a proteome-wide scale^9^, and is an imperative but still challenging component to modern glycoproteomic studies^10^.

Currently, liquid chromatography coupled with tandem mass spectrometry (LC-MS/MS) is the method of choice widely used in proteomics and glycoproteomics^11,12^. Novel MS/MS fragmentation methods derived from higher-energy collisional dissociation (HCD) and electron transfer dissociation (ETD), such as stepped collision energy HCD (SCE-HCD)^13^ and ETD with supplemental HCD (EThcD)^14^, have been proven powerful for intact glycopeptides profiling. The most common strategy for peptide identification uses the data-dependent acquisition (DDA) approach, in which MS/MS (MS2) fragmentation is triggered by precursor ions observed in a full mass range survey scan (MS1). Various software tools^15^, such as pGlyco 2.0^16^, MSFragger-Glyco^17^ and O-Pair Search^18^, have been developed for the interpretation of DDA data of intact glycopeptides. However, a major bottleneck of the DDA approach is that the precursor selection constitutes a stochastic element^19^, resulting in the “missing value” problem.

To overcome this limitation, data-independent acquisition (DIA) methods have been proposed^20–22^, including a representative variant named sequential window acquisition of all theoretical mass spectra (SWATH-MS)^23^, where the instrument acquires fragmentation information of all precursor ions within defined isolation windows in a systematic manner. DIA has been reported to achieve deep proteome coverage capabilities with quantitative consistency and accuracy for large-scale proteomic studies^24^, and is now starting to be applied to the field of glycoproteomics^25^. Based on standard DIA protocols developed for proteomics, DIA methods have been established for targeted analysis of intact N-glycopeptides^26–29^, in which spectral libraries were built by adding manually curated glycan fragment Y ions. Zhou et al. developed a SWATH-MS method with optimized variable windows for a set of target glycopeptides to allow accurate glycoform measurement^30^. These methods have achieved better sensitivity than DDA for analyzing glycoforms of several or a dozen of glycoproteins. In 2019, Ye et al. proposed Glyco-DIA, a DIA-based strategy for O-glycoproteomics, enabling high-throughput quantitative O-GalNAc type glycoproteomic analysis in complex biological samples^31^.

Error rate control for glycopeptide identification is essential but particularly complicated in DIA analyses due to the increased complexity of DIA MS/MS spectra originated from multiple co-eluted precursors, especially when using wide isolation windows. In the case of HCD MS/MS, the same set of fragment ions could be generated for glycopeptides common in peptide sequence but different in glycan, resulting in a high level of misinterpretations of DIA data^25^. Although a few studies have elucidated error rate estimation for DDA-based glycopeptide analyses^16,32–34^, statistical control of DIA-based proteome-wide glycopeptides analyses, to the best of our knowledge, has not been properly addressed.

Herein, we propose GproDIA, a pipeline that applies the concept of peptide-centric DIA analysis to proteome-wide characterization of intact glycopeptides. GproDIA provides comprehensive statistical control by a 2-dimentional false discovery rate (FDR) approach and a glycoform inference algorithm, enabling accurate glycopeptide identification using wide isolation windows. We further adapt a semi-empirical spectrum prediction strategy to expand the coverage of spectral libraries for glycopeptides. We benchmark GproDIA for N-glycopeptide profiling on DIA data of yeast and human serum samples, demonstrating that DIA with GproDIA outperforms DDA in terms of capacity and data completeness of identification, as well as accuracy and precision of quantification.

## Results

### GproDIA enables characterization of intact glycopeptides from DIA data

GproDIA was developed based on the principle of peptide-centric analysis, which has been commonly used for detection of peptides from DIA data^24^. The workflow is presented in **Fig. 1**. First, a spectral library of glycopeptides is built by DDA with pre-fractionation or using a long LC gradient. As LC conditions are different from those used for non-glycosylated peptides, instead of using iRT^35^ as exogenous standards for retention time (RT) normalization, an extra DDA injection of the glycopeptides sample is performed with the same LC condition as that used for DIA experiments. The shared identifications between different LC conditions are used as internal standards to calibrate library RT to the gradient used in DIA. An example of RT calibration is shown in **Supplementary Fig. 1.** The spectral library contains the RT of glycopeptides, the precursor *m*/*z*, and *m*/*z* and intensities of annotated fragments in SCE-HCD MS/MS including peptide fragments (b/y, with or without one HexNAc residue and its cross-ring fragment) and intact peptide with glycan fragments (Y)(**Fig. 1a**). Next, three types of decoy libraries are generated by adding random mass shifts to glycan fragment peaks^36^ (glycan decoy), reversing the peptide sequences (peptide decoy), and performing the two operations successively (both decoy) (**Fig. 1b**).

**Fig. 1.**
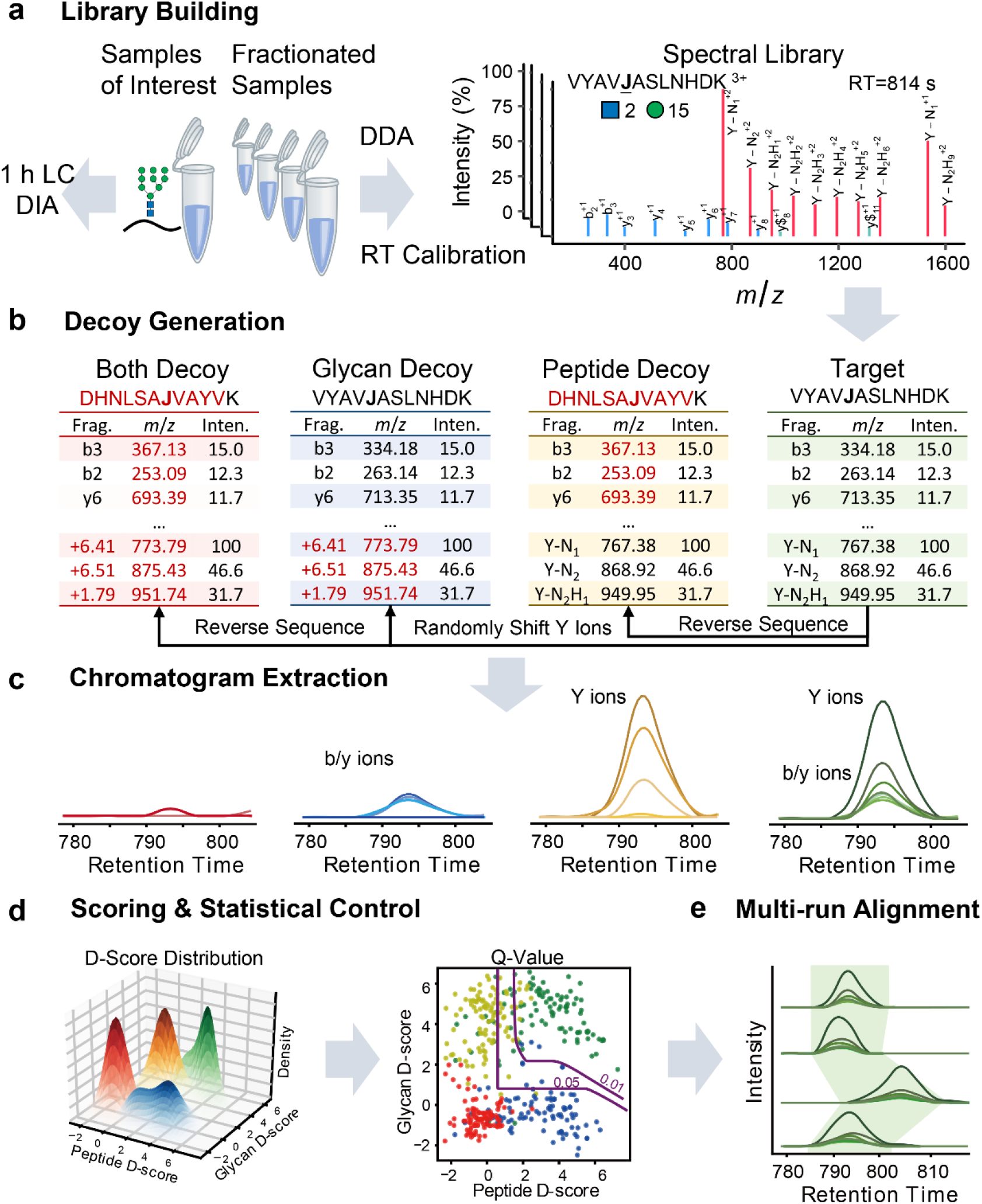
The workflow of GproDIA. (**a**) Building a spectral library of glycopeptides containing peptide ions (blue lines for b/y, and green lines for b/y with one HexNAc or its cross-ring fragment) and glycan Y ions (red lines) by DDA. “J” in peptide sequence indicates the N-glycosylation site. The glycan symbols are as follows: a green circle or “H” represents Hex; a blue square or “N” represents HexNAc. (**b**) Generating peptide decoys, glycan decoys and both decoys. (**c**) Extracting chromatogram features from the DIA data. (**d**) Scoring the extracted features and estimating error rates by a 2-dimentional FDR approach. (**e**) Performing multi-run alignment to reduce missing values. (**c-e**) Green color indicates target peak groups, yellow indicates peptide decoy peak groups, blue indicates glycan decoy peak groups, and red indicates both decoy peak groups.

Then OpenSWATH^37^ is used to extract chromatogram data of the target glycopeptides and the decoys from the DIA data (**Fig. 1c**), and the extracted peak group features are scored using a semi-supervised learning approach implemented by PyProphet^38^. A 2-dimentional FDR approach is used to estimate error rates of identification (**Fig. 1d**). In brief, the distributions of discriminant scores (D-scores) of the targets and decoys are fitted using a bivariate four-groups mixture model. Local FDR (namely posterior error probability, PEP) of each target peak group is computed from D-score density (**Supplementary Fig. 2**). Global FDR (q-value) is then derived from PEPs. Finally, TRIC^39^ is used for multi-run alignment to reduce missing values (**Fig. 1e**).

### Benchmarking using data from yeast samples

For benchmark purposes, we performed DDA and DIA experiments using an 1 h LC gradient with 4 technical replicates, as well as a DDA using a 6 h LC gradient with 3 technical replicates, on a sample of fission yeast (*Schizosaccharomyces pombe*) glycopeptides. pGlyco 2.0^16^ was used for database searching of the DDA data, and 1% GPSM-level FDR cutoff was used. A sample-specific spectral library (fission yeast SSL, **Supplementary Table 1**) was built using the 6 h DDA data for DIA data analysis by GproDIA. For DIA, results with q-value < 5% in each run and q-value < 1% in at least one run at peak group level, as well as q-value < 1% at glycopeptide level in the global context^38^, were reported. Statistics of the results are shown in **Fig. 2** and **Supplementary Fig. 3-5**. In average, 418 ± 2 (mean ± standard deviation, sic passim) glycopeptide precursors corresponding to 348 ± 2 site-specific glycans were detected per replicate run from the DIA data (**Fig. 2a** and **Supplementary Fig. 5a**), more than those identified from the 1 h DDA (357 ± 16 precursors and 293 ± 9 site-specific glycans, Supplementary Fig. 3a) and the 6 h DDA (351 ± 12 precursors and 289 ± 9 site-specific glycans, **Supplementary Fig. 4a**). Notably, we use the term “site-specific glycan” referring glycans on specific glycosylation sites in a group of glycoproteins that are not distinguishable by protein inference^16^, rather than positions of glycans on peptide sequences. Indeed, it is not very common that an N-glycopeptide has more than one potential glycosylation sites.

**Fig. 2.**
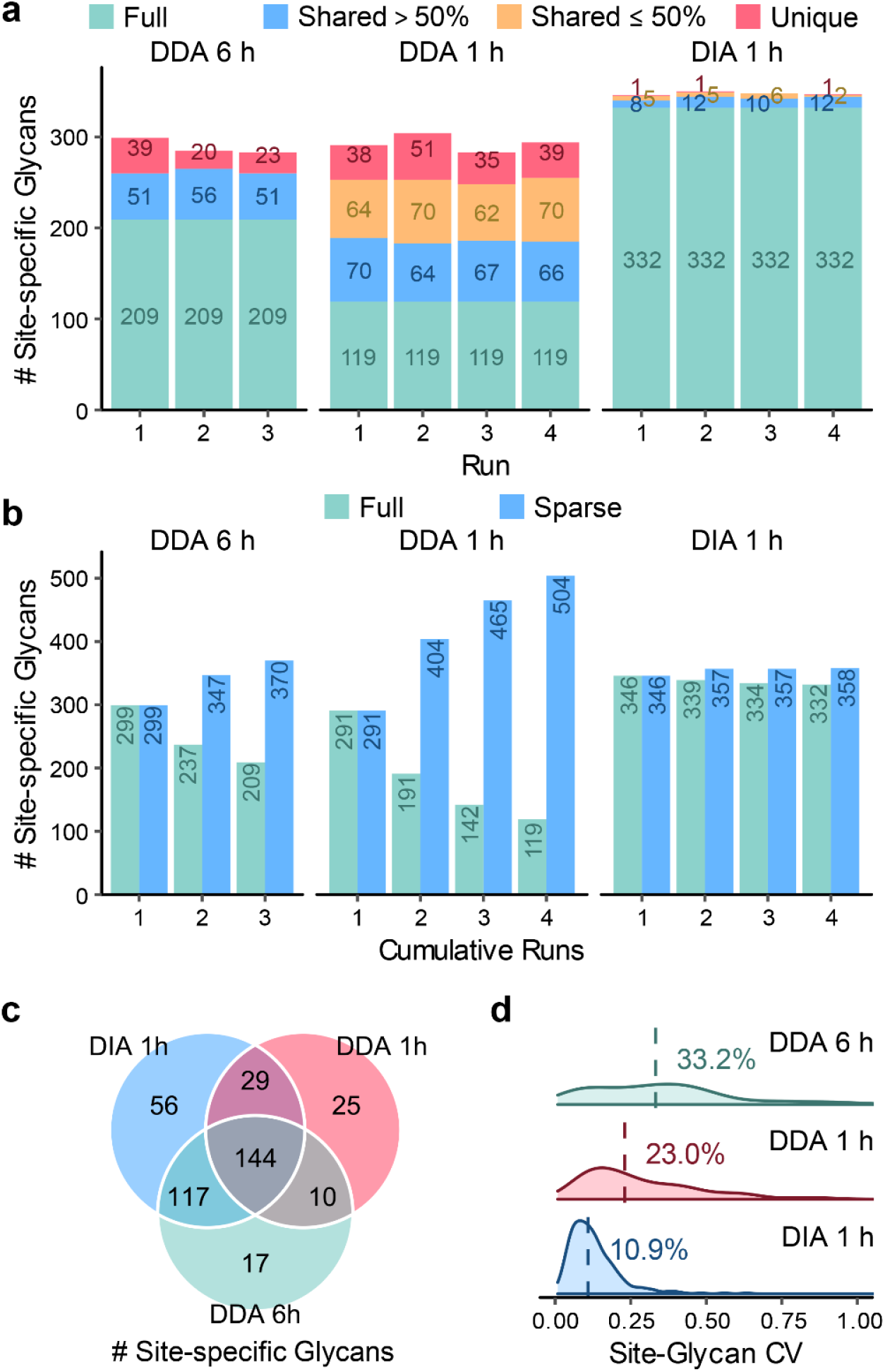
Performance comparison of DDA and DIA on the fission yeast sample at the site-specific glycan level. (**a**) Numbers of identifications per run. “Full” represents identifications observed in all the runs; “shared >50%” represents identifications observed in >50% runs; “shared ≤50%” represents identifications observed in >1 but ≤50% runs; “unique” represents identifications observed in only 1 run. (**b**) Numbers of cumulative identifications across runs. “Full” represents identifications shared in the cumulative runs; “sparse” represents identifications observed in at least one run in the cumulative runs. (**c**) Comparison of numbers of identifications shared in >50% runs using DDA with an 1 h LC gradient, DDA with a 6 h LC gradient, and DIA with an 1 h LC gradient. (**d**) Coefficients of variation (CVs) of quantification results. Medians are indicated.

From the 4 DIA replicate runs, 433 glycopeptide precursors corresponding to 358 site-specific glycans on 142 protein glycosites were detected totally. Among them, 91% (392) precursors, 93% (332) site-specific glycans, and 96% (136) glycosites were shared in all the replicates (**Fig. 2b** and **Supplementary Fig. 5b**), indicating much fewer missing values than those of 1h DDA (122/666 = 18% at precursor level, 119/504 = 24% at site-specific glycan level, and 98/190 = 52% at glycosite level, **Supplementary Fig. 3b**) and 6 h DDA (246/461 = 53% at precursor level, 209/370 = 56% at site-specific glycan level, and 102/139 = 73% at glycosite level, **Supplementary Fig. 4b**). Considering identifications shared in >50% replicate runs, DIA detected 21% more (418/346) glycopeptide precursors, 20% more (346/288) site-specific glycans, and 19% more (139/117) protein glycosites than 6 h DDA, as well as 84% more (418/227) precursors, 66% more (346/208) site-specific glycans, and 14% more (139/122) protein glycosites than 1 h DDA (**Fig. 2c** and **Supplementary Fig. 5c**). It should be noticed that a less strict error rate control was applied on DDA results (only a GPSM-level FDR cutoff) than that on DIA results (peak group q-value and global glycopeptide q-value). Coefficients of variation (CVs) of glycopeptide precursor, site-specific glycan and protein glycosite quantification results were calculated among the technical replicates as shown in **Fig. 2d** and **Supplementary Fig. 3c**, **4c** and **5d**. The median CVs were ~11% using DIA, much smaller than those using 1 h DIA (~17% at precursor level, ~23% at site-specific glycan level, and ~37% at protein glycosite level) and 6 h DDA (>32%). We also present the DIA results without multi-run alignment (1% peak group q-value and 1% global glycopeptide q-value) in **Supplementary Fig. 6**, wherein glycopeptides identification and quantification results close to the ones with multi-run alignment were obtained. The results indicate that the DIA workflow using GproDIA outperforms DDA not due to the multi-run alignment, but originated from the inherent feature of systematic and panoramic MS/MS recording in DIA that provides broadly informative data.

As an alternative to sample-specific libraries, community spectral libraries such as Pan-Human^40^ can be effectively used for peptide-centric DIA data analysis^41^. We tested the feasibility of using a lab repository-scale spectral library (fission yeast LRL, **Supplementary Table 1**) generated by combining the SSL library and fission yeast data of previous projects in our labs. The results are presented in **Supplementary Fig. 7**. Using LRL, 18% more (495/418) glycopeptide precursors, 14% more (394/346) site-specific glycans, and 3% more (143/139) protein glycosites were detected in >50% replicate runs than using SSL, while the CVs (~11%) were very close to those using SSL. DIA analysis was also performed on a budding yeast (*Saccharomyces cerevisiae*) sample with 3 technical replicates in an 1 h LC gradient, using a spectral library built from budding yeast DDA data (**Supplementary Table 1**). Low levels of missing values and CVs were also observed (**Supplementary Fig. 8**). All the results suggest that, within the tested condition, DIA with GproDIA improves the number of identifications and reproducibility of glycopeptide characterization compared to DDA-based workflows.

### Inference of glycoforms in wide isolation windows

We further compared DDA and DIA on a human serum sample acquired with 3 technical replicates using the 1 h LC gradient (**Supplementary Fig. 9** and **10**). The DIA data were analyzed using a sample-specific spectral library built by DDA with pre-fractionation (serum SSL, Supplementary Table 1). Unlike high-mannose-type glycans of yeast, glycans in human serum have more complex compositions, presenting greater challenges for DIA analysis. For yeast samples, glycopeptides with the same peptide sequence and different glycans (HexNAc_2_Hex_*n*_) have mass differences of at least one hexose residue (162 Da), and are hardly co-fragmented in an isolation window (25 *m*/*z* in this study). For human serum samples, however, DIA analysis suffers from potential interference of glycopeptides with the same peptide sequence but different glycans (referred to as “glycoforms”) in the same isolation window. Therefore, although a large number of glycopeptides were reported by DIA (**Supplementary Fig. 10**), the results can have high error rates of identification in the glycan part.

Inspired by IPF^42^, a DIA data analysis tool for peptides carrying PTMs, we further developed an algorithm to evaluate the global FDR (q-value) at glycoform level. The workflow is illustrated in **Fig. 3a-c** and **Supplementary Fig. 11**. In brief, all theoretical Y fragment ions (called “identification transitions”) are generated in silico for each target glycopeptide precursor and the corresponding potential glycoforms within the isolation window when building the spectral library. The potential glycoforms were collected from the pGlyco 2.0 built-in glycan database^16^, containing 3065 glycan structures, and might not present in the original library. Signals of precursors of target glycopeptides and identification transitions of the target glycopeptides/glycoforms are traced during chromatogram extraction from DIA data. The PEP of MS2 peak groups (PEP_MS2_), precursors (PEP_MS1_) and identification transitions (PEP_transition_) are integrated using a Bayesian hierarchical model (BHM), leading to a glycoform-level posterior probability (PP) for each detected peak group, from which the global FDR (q-value) at glycoform level can be derived.

**Fig. 3.**
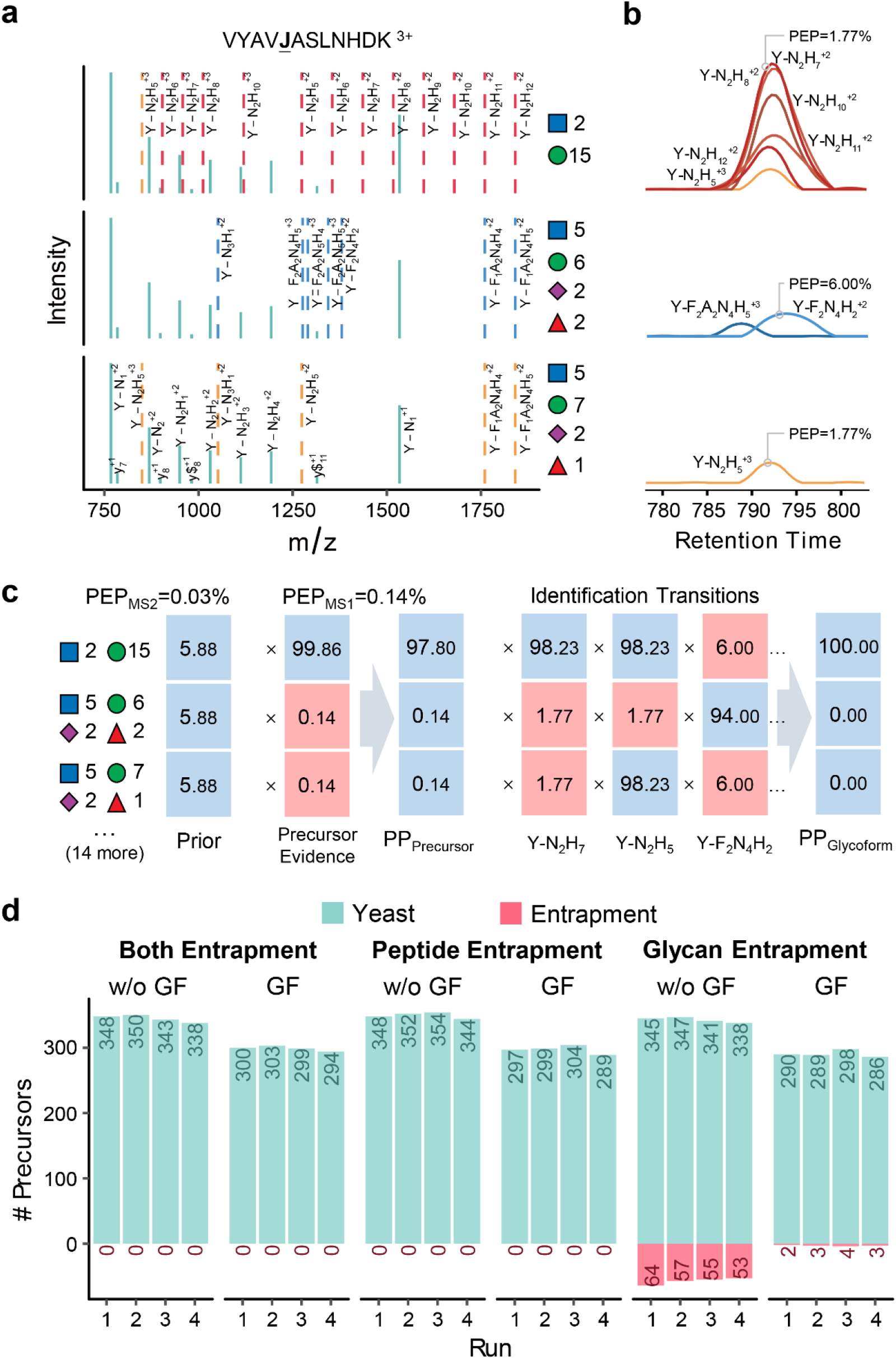
Inference of glycoforms. (**a**) Generating theoretical Y fragment ions as identification transitions (dashed lines) for the target glycopeptide and all potential glycoforms. “J” in peptide sequence indicates the N-glycosylation site. The glycan symbols are as follows: a green circle or “H” represents Hex; a blue square or “N” represents HexNAc; a red triangle or “F” represents Fuc; a purple diamond or “A” represents NeuAc. (**b**) Extracting chromatograms of identification transitions from the DIA data and performing transition-level scoring to get the PEPs of transitions. (**c**) Integrating the precursor and transition PPs to glycoform PPs using a Bayesian hierarchical model. (**d**) Numbers of identifications without (w/o GF) or with (GF) glycoform inference of the fission yeast sample using the entrapment libraries.

The performance of the glycoform-level FDR control was tested on the fission yeast data using an entrapment strategy by adding glycopeptides with peptide sequences (peptide entrapment) or glycans (glycan entrapment) or both (both entrapment) from human serum SSL to the fission yeast SSL library. In all the analyses, we kept the number of entrapment glycopeptide precursors similar to that of the yeast library (**Supplementary Table 1**). Since we focus on investigating the performance of the glycoform-level error rate control, multi-run alignment was not performed, and no global glycopeptide-level q-value filter was applied. The DIA analyses results are presented in **Fig. 3d**. Using the entrapment library containing human glycopeptides (both entrapment), fewer identifications were observed compared to those using the fission yeast library, because the entrapment library contains a large fraction of “false targets” that are not detectable in the sample (referred to as π_0_^38^), compromising the detection sensitivity. Nevertheless, no entrapment identifications were observed at 1% peak group-level q-value. Similar results were obtained using the entrapment library containing glycopeptides with peptide sequences from human and glycans from yeast (peptide entrapment), suggesting satisfactory performance of error rate control in the peptide part. Using the entrapment library containing glycopeptides with peptide sequences from yeast and glycans from human (glycan entrapment), although 1% peak group-level q-value filter was applied, there were in average 14% of entrapment identifications (to all of those identified) remaining per run without glycoform inference. After applying 1% glycoform-level q-value filter, despite a loss of 15% yeast identifications due to poor signals of precursors and/or glycoform-specific fragments, the entrapment percentage declined to ~1%.

### Benchmarking using data from human serum and mixed-organism samples

GproDIA was then tested on the human serum data with glycoform inference enabled. In addition to the sample-specific library (serum SSL), a lab repository-scale spectral library (serum LRL, **Supplementary Table 1**) was also used for DIA data analysis. Global glycopeptide-level q-value cutoff was 1%. After multi-run alignment, results with glycoform-level q-value < 5% in each run and < 1% in at least one run were finally reported (**Supplementary Fig. 12** and **13**). Comparison between DDA and DIA results is illustrated in **Fig. 4a-d**. At site-specific glycan level, compared to DDA, DIA using SSL and LRL brought 14% more (539/474) and 35% more (638/474) identifications, respectively, in average per run, whereas fewer missing values (463 shared in all replicates/559 in total using SSL, and 531/733 using LRL) by DIA were observed than DDA (262/717). Considering identifications shared in >50% replicate runs, DIA using SSL and LRL detected 26% more (556/443) and 47% more (650/443) site-specific glycans, respectively, than DDA. CVs using DIA (12.8% with SSL and 12.2% with LRL) were significantly smaller than that using DDA (26.1%).

**Fig. 4.**
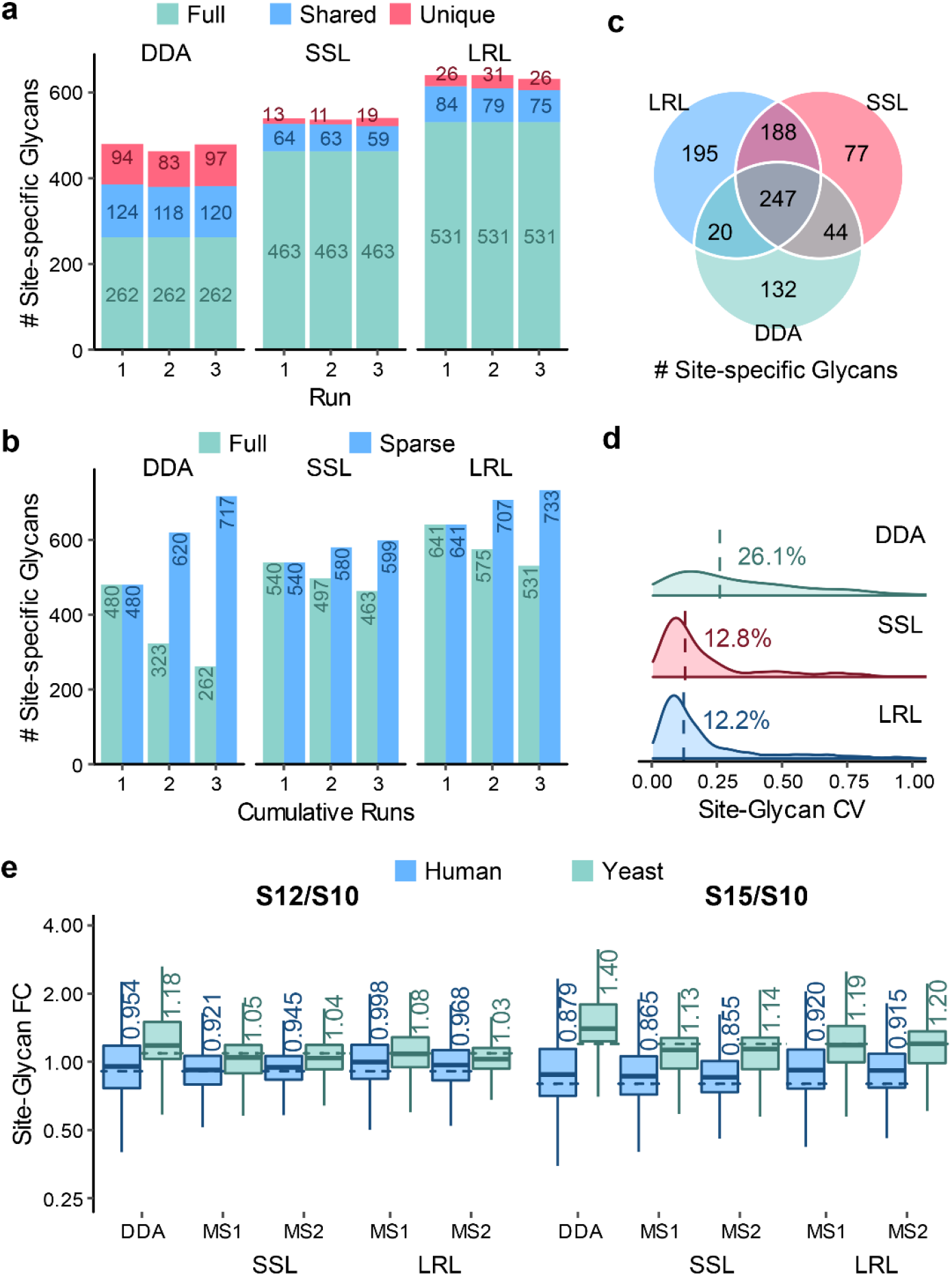
Performance comparison of DDA and DIA on the human serum and mixed-organism samples at the site-specific glycan level. (**a**) Numbers of identifications from the serum sample per run. “Full” represents identifications observed in all the runs; “shared” represents identifications observed in 2 runs; “unique” represents identifications observed in only 1 run. (**b**) Numbers of cumulative identifications from the serum sample across runs. “Full” represents identifications shared in the cumulative runs; “sparse” represents identifications observed in at least one run in the cumulative runs. (**c**) Comparison of numbers of identifications from the serum sample shared in >50% runs using DDA, DIA with the sample-specific library (SSL), and DIA with the lab repository-scale library (LRL). (**d**) Coefficients of variation (CVs) of quantification results. Medians are indicated. (**e**) Box plot visualization of fold change of the quantification results of the mixed-organism samples. Percent changes were calculated based on the mean quantities in three replicates of each sample. The medians are indicated. The boxes indicate the interquartile ranges (IQR), and whiskers indicate 1.5 × IQR values; no outliers are shown. The dashed lines indicate theoretical fold changes of the organisms (1:0.9:0.8 (S10:S12:S15) for human and 1:1.1:1.2 (S10:S12:S15) for yeast).

The performance of GproDIA was further evaluated on data of mixed-organism samples containing glycopeptides from budding yeast and human serum with different abundance (S10, S12, and S15). DIA analysis was performed using a combined library of the budding yeast library and the serum SSL library, as well as a combined library of the budding yeast library and the serum LRL library, respectively (**Supplementary Table 1** and **Supplementary Fig. 14-16**). Based on the mean quantities in three replicates of each sample, fold changes of detected site-specific glycans of samples S12/S10 and S15/S10 were calculated and visualized in **Fig. 4e**. Fold changes of human and yeast glycopeptide abundance were closer to the theoretical values using DIA-based quantification at both MS1 and MS2 level with the yeast + serum SSL library than those using DDA. Using the yeast + serum LRL library, fold changes of human glycopeptide abundance were overestimated, while fold changes of yeast glycopeptides were measured accurately. In addition, the distribution of fold changes was less dispersed using DIA than that using DDA. All the results underline the high quality of DIA-based glycopeptide characterization using GproDIA with glycoform inference that outperforms DDA-based workflows in terms of numbers of identifications, as well as accuracy and precision of quantification.

### Extending library coverage semi-empirically

In peptide-centric DIA data analysis, the capability of detection is limited due to the incomplete coverage of spectral libraries. For this reason, we propose a computational approach to expand the coverage of spectral libraries of glycopeptides semi-empirically, wherein the MS2 spectra of predicted glycopeptides are generated by swapping and combining the peptide and glycan fragment peaks in experimental spectra of different glycopeptides using a *k*-nearest neighbor (KNN) strategy (**Fig. 5a**). The RT of the predicted glycopeptides are the weighted mean RT of experimentally identified glycopeptides with the same peptide sequences and close monosaccharide compositions by the KNN strategy.

**Fig. 5.**
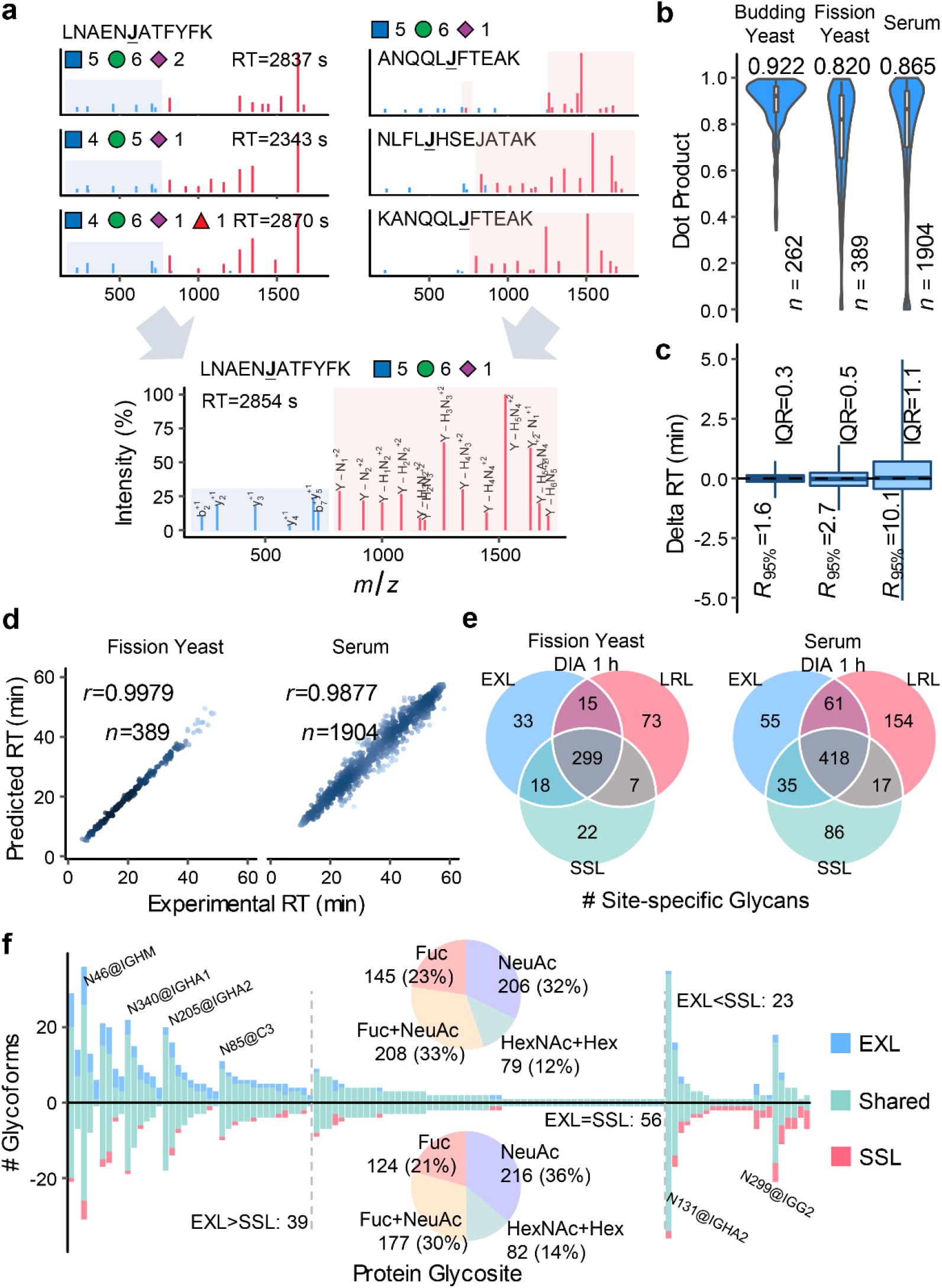
Generating and utilizing semi-empirical spectral libraries. (**a**) Generating semi-empirical MS/MS spectra by combining the peptide and glycan fragment peaks in the experimental spectra of different glycopeptides. “J” in peptide sequence indicates the N-glycosylation site. The glycan symbols are as follows: a green circle or “H” represents Hex; a blue square or “N” represents HexNAc; a red triangle or “F” represents Fuc; a purple diamond or “A” represents NeuAc. (**b**) The distributions of dot products computed between the predicted and experimental MS/MS peak intensities. (**c**) The differences between predicted and experimental retention times (RTs). The medians are indicated. The boxes indicate the interquartile ranges (IQR), and the whiskers show the ranges between 2.5% and 97.5% percentiles (*R*_95%_); no outliers are shown. (**d**) Pearson correlation coefficients (*r*) between predicted and experimental RTs. “*n*” indicates the number of generated MS2 spectra or RT values by the prediction method. (**e**) Comparison of numbers of identifications shared in >50% runs using the sample-specific library (SSL), the lab repository-scale library (LRL), and the extended library by the semi-empirical approach (EXL). (**f**) Comparison of numbers of glycoforms per glycosite detected from serum using SSL and EXL. Proportions of glycan composition types are shown in the pie charts. Glycosites with high variabilities in glycoforms are indicated on the figure.

Cross validations of the prediction were conducted using the fission yeast LRL library and the budding yeast library, wherein the library entry of each glycopeptide precursor was pulled out from the library, and meanwhile the rest of the library entries were used to generate the predicted library entry of the glycopeptide precursor. Different numbers of nearest neighbors (*k*) were tested. Dot products (DPs) were computed between the predicted and experimental MS/MS intensities (**Supplementary Fig. 17a** and **18a**). With *k* = 3, the median DP was 0.820 for the fission yeast library and 0.922 for the budding yeast library, higher than those without KNN and with *k* = 2 or 4. Increasing *k* can lead to higher prediction accuracy, but fewer glycopeptides can be predicted because at least *k* neighboring entries for generating the peptide part and the glycan part are required in the experimental library. Therefore, we chose *k* = 3 to achieve the trade-off between prediction accuracy and library coverage. With *k* = 3, the interquartile ranges (IQRs) of the differences between predicted and experimental RTs were <0.6 min (**Supplementary Fig. 17b** and **18b**). Pearson correlation coefficients (*r*) of predicted and experimental RTs were >0.99 (**Fig. 5d**, **Supplementary Fig. 17c** and **18c**). The serum SSL library was also used for the cross validation with *k* = 3. The median DP was 0.865 (**Fig. 5b**), the IQR of RT differences was 1.1 min (**Fig. 5c**), and the r of RTs was >0.98 (**Fig. 5d**). The prediction on human serum glycopeptides is less accurate compared to yeast due to the high complexity of human serum glycopeptides. DPs, RT differences and *r* were also computed among replicate experimental spectra, which can be considered as possible upper limit of prediction accuracy (**Supplementary Fig. 19**), showing that there are still rooms for improvement. Nevertheless, due to the limit of current available data size of glycopeptides and the highly complex isomeric glycan structures, more accurate prediction by machine learning and deep learning strategies can hardly be applied at this stage.

Next, we built a semi-empirical library based on the fission yeast SSL library, which was then merged with the SSL to form an extended library (fission yeast EXL, **Supplementary Table 1**), containing 331 extra precursors and 288 site-specific glycans that are not present in the SSL (**Supplementary Fig. 20a**). From the fission yeast DIA data, 5% more (365/346) site-specific glycans were detected in >50% runs using EXL than those using SSL, while the CVs stayed unchanged approximately (**Fig. 5e** and **Supplementary Fig. 21**). Performance of error rate control when using extended libraries was also evaluated using the entrapment strategy (**Supplementary Fig. 22**).

An extended library (serum EXL, **Supplementary Table 1**) was also generated from the serum SSL library. To avoid combinatorial explosion of peptides and glycans, we collected a list of glycopeptides combined from the serum LRL library and a publication on N-linked intact glycopeptides in human serum^43^, and only peptide-glycan combinations in the glycopeptide list were taken into consideration when generating the extended library. Consequently, the EXL library containing 1990 extra precursors and 927 site-specific glycans that are not present in the serum SSL (**Supplementary Fig. 20b**). Combining all the 3 replicates, 118 protein glycosites were detected in total using both the SSL and EXL libraries. Among them, on 39 glycosites, more glycoforms were detected using EXL than using SSL, while on 23 glycosites, more glycoforms were detected using SSL (**Fig. 5f**). The increase in the detected glycoforms can be attributed to the greater proportion of fucosylated glycans (145 site-specific glycans with fucosylation and without sialylation, as well as 208 with both fucosylation and sialylation) detected using EXL than using SSL (124 with fucosylation and without sialylation, as well as 177 with both fucosylation and sialylation). It should be noted that the semi-empirical approach does not increase the protein glycosites. The CVs stayed unchanged approximately using the human serum EXL (**Supplementary Fig. 23**).

### Discussions

With more efficient usage of ions, DIA can provide a significant increase in the identification efficiency of glycopeptides compared to DDA^25,44^. We have benchmarked our DIA-based workflow against the DDA-based glycoproteomics, demonstrating that short-gradient DIA can outperform DDA under the same conditions or even with a much longer gradient in terms of detectable glycopeptides as well as measurement reproducibility. It has been reported that DIA copes well with shortening of the LC gradient length because the deterministic nature of the MS2 sampling in DIA attenuates the attrition in number of identifications for shorter separation gradients, while the number of acquired MS/MS spectra and identifications decrease proportionally with the gradient length in DDA mode^24^. As a less time-consuming approach for intact glycopeptides profiling, short-gradient DIA is favorable for large-scale quantitative glycoproteomic analyses.

Interference from other co-eluted and co-fragmented glycopeptides is the main challenge for DIA data analysis that may result in a high level of misinterpretations. In GproDIA, we adapt a 2-dimensional FDR approach and a glycoform inference strategy, providing comprehensive statistical control for glycopeptide identification. Glycoform inference can be used as an option to perform more strict assessment by utilizing signals of precursors and glycan-specific fragments to resolve interference from potential glycoforms. In this study, glycoform inference was enabled when multiple glycopeptide precursors with the same peptide sequence and different glycans are arranged to be fragmented in one isolation window. Despite the herein proposed strategies to control error rates when using wide isolation windows, we still recommend to design the isolation windows properly according to the mass distribution of glycopeptides^30^, if possible, for improving the detection sensitivity. Recent advances in ion mobility spectrometry (IMS) including high field asymmetric waveform ion mobility spectrometry (FAIMS)^45^ and parallel accumulation-serial fragmentation (diaPASEF)^46^ have achieved rapid improvements in the sensitivity of DIA analysis. We expect that DIA-based glycopeptides profiling can benefit from the enhanced separation of glycoforms by IMS.

Spectral library coverage determines the upper limit of the identification capacity by peptide-centric DIA analysis. To date, the majority of DIA studies have used DDA-based sample-specific spectral libraries, frequently with pre-fractionation (for the serum sample in this study), or sometimes by repeated DDA analysis of non-fractionated samples (for the fission yeast sample)^24^. Other sources can also be considered as a supplement for library completeness. We achieve the best coverage by using repository-scale libraries integrating data from multiple previous projects in our lab. We envision that more “off-the-shelf” glycopeptide libraries will be built and published by the community, just like those for proteomic studies^40^, which can then be used for glycopeptide DIA data analysis.

Since repository-scale libraries may not always be available, we have built semi-empirical libraries as an attempt to extend the library coverage. In such predicted libraries, it is common that a significant fraction of glycopeptides are actually not present in the samples at a detectable level, and the large query space may increase the multiple testing burden and compromise detection sensitivity^24^. Instead of enumerating all the peptide-glycan combinations in a spectral library, we suggest researchers focus on a subset of glycoproteins/glycoproteoforms of interest for their specific biological questions. The current proposed strategy facilitates detection of more glycoforms by DIA on protein glycosites that are observed by DDA, but cannot increase the coverage at glycosite level. However, deep learning-based tools such as Prosit^47^ and DeepDIA^48^ have been developed for generating in silico peptide spectral libraries directly from peptide sequences, and we anticipate that predicted libraries will break through the limitation on coverage of glycopeptide libraries by DDA in the future when large scale glycopeptide libraries are built by the community.

Although it is demonstrated here in the context of N-glycoproteomics, the principle of comprehensive statistical control in GproDIA is also applicable to O-glycopeptides for peptide sequence and glycan identification. O-glycopeptides generally have multiple serine and/or threonine residues that serve as potential glycosites. Unfortunately, HCD is not sufficient for glycosite localization for O-glycopeptides^10^, and to date, HCD is the only method of choice for DIA. We hope that further instrumental development implements ETD-based DIA methods for site-specific O-glycopeptide analyses.

## Code availability

A test version of GproDIA is available on request to the correspondence author at.

