## Supplementary Information for "GproDIA enables data-independent acquisition glycoproteomics with comprehensive statistical control"

Yi Yang et al.

**Supplementary Table 1.** Spectral libraries used in this study.

| Name | Description |
|---|---|
| Fission yeast SSL | A sample-specific spectral library of fission yeast generated from DDA data using a 6 h LC gradient with 3 replicates. An extra DDA injection with an 1 h LC gradient has been used for RT calibration. Data with the 1 h gradient have also been appended to the calibrated library.<br>Containing 502 precursors of 434 glycopeptides, 412 site-specific glycans, 156 protein glycosites (excluding decoys). |
| Fission yeast LRL | A lab repository-scale spectral library of fission yeast generated by combining the SSL library and fission yeast data of previous projects in our labs [a]. RTs have been calibrated to the 1 h LC gradient.<br>Containing 1044 precursors of 832 glycopeptides, 725 site-specific glycans, 235 protein glycosites (excluding decoys). |
| Fission yeast EXL | An extended spectral library of fission yeast generated by combining the SSL library and a semi-empirical library generated from the SSL library.<br>Containing 833 precursors of 733 glycopeptides, 700 site-specific glycans, 156 protein glycosites (excluding decoys). |
| Budding yeast | A lab repository-scale spectral library of budding yeast generated from budding yeast DDA data. RTs have been calibrated to the 1 h LC gradient using an extra injection of DDA.<br>Containing 850 precursors of 667 glycopeptides, 613 site-specific glycans, 241 protein glycosites (excluding decoys). |
| Serum SSL | A sample-specific spectral library of human serum generated from DDA data using an 1 h LC gradient with 20 fractions. An extra DDA injection with an 1 h LC gradient without fractionation has been used for RT calibration. Data with the 1 h gradient and without fractionation have also been appended to the calibrated library.<br>Containing 3518 precursors of 2402 glycopeptides, 2082 site-specific glycans, 396 protein glycosites (excluding decoys). |
| Serum LRL | A lab repository-scale spectral library of human serum generated by combining the SSL library and serum data of previous projects in our labs [a]. RTs have been calibrated to the 1 h LC gradient.<br>Containing 5734 precursors of 4011 glycopeptides, 3519 site-specific glycans, 571 protein glycosites (excluding decoys). |
| Serum EXL | An extended spectral library of human serum generated by combining the SSL library and a semi-empirical library generated from the SSL library.<br>Containing 5508 precursors of 3433 glycopeptides, 3009 site-specific glycans, 396 protein glycosites (excluding decoys). |
| Yeast + serum SSL | A spectral library generated by combining the budding yeast library and the serum SSL library. |
| Yeast + serum LRL | A spectral library generated by combining the budding yeast library and the serum LRL library. |
| Both entrapment | A spectral library generated by combining the fission yeast SSL library and 500 precursors randomly sampled from the serum SSL library. |

| | |
|---|---|
| Peptide entrapment | A spectral library generated by combining the fission yeast SSL library and 500 entrapment precursors [b]. The entrapment precursors have been generated semi-empirically using the fission yeast and serum SSL libraries, and then randomly subsampled to 500. The peptide sequences of the entrapment precursors are from human, while the glycans are from yeast. |
| Glycan entrapment | A spectral library generated by combining the fission yeast SSL library and 500 entrapment precursors [b]. The entrapment precursors have been generated semi-empirically using the fission yeast and serum SSL libraries, and then randomly subsampled to 500. The peptide sequences of the entrapment precursors are from yeast, while the glycans are from human. |
| Both entrapment EXL | A spectral library generated by combining the fission yeast EXL library and 850 precursors randomly sampled from the serum SSL library. |
| Peptide entrapment EXL | A spectral library generated by combining the fission yeast EXL library and 850 entrapment precursors [b]. The entrapment precursors have been generated semi-empirically using the fission yeast and serum SSL libraries, and then randomly subsampled to 850. The peptide sequences of the entrapment precursors are from human, while the glycans are from yeast. |
| Glycan entrapment EXL | A spectral library generated by combining the fission yeast EXL library and 850 entrapment precursors [b]. The entrapment precursors have been generated semi-empirically using the fission yeast and serum SSL libraries, and then randomly subsampled to 850. The peptide sequences of the entrapment precursors are from yeast, while the glycans are from human. |

[a] The combination of libraries is performed using the approach of generating "consensus" spectrum when multiple spectra for a glycopeptide exist in different libraries. During the procedure, some glycopeptides can be eliminated if the consensus spectrum doesn't meet the criteria of transition selection.

[b] As a special case, when generating entrapment libraries using the semi-empirical approach, variants with the same glycan monosaccharide compositions but different isomeric glycan structures are regarded as different precursors which can have different glycan fragment peaks. The same criteria are followed when counting entrapment identifications in DIA results.

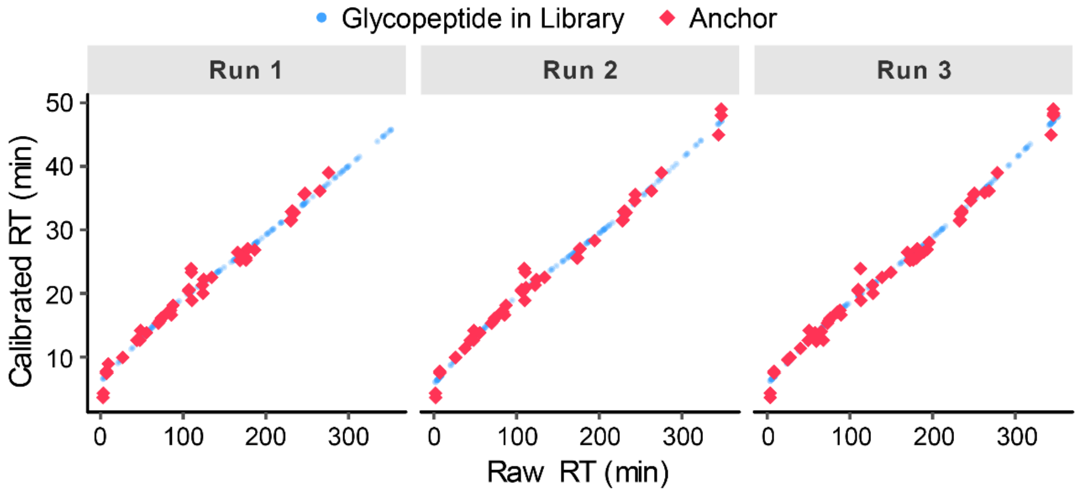


**Supplementary Fig. 1.** RT calibration for the fission yeast SSL library. RTs of glycopeptides in a 6 h LC gradient were transformed into an 1 h LC gradient. The shared identifications between the 6 h and the 1 h LC gradients were used as anchors.

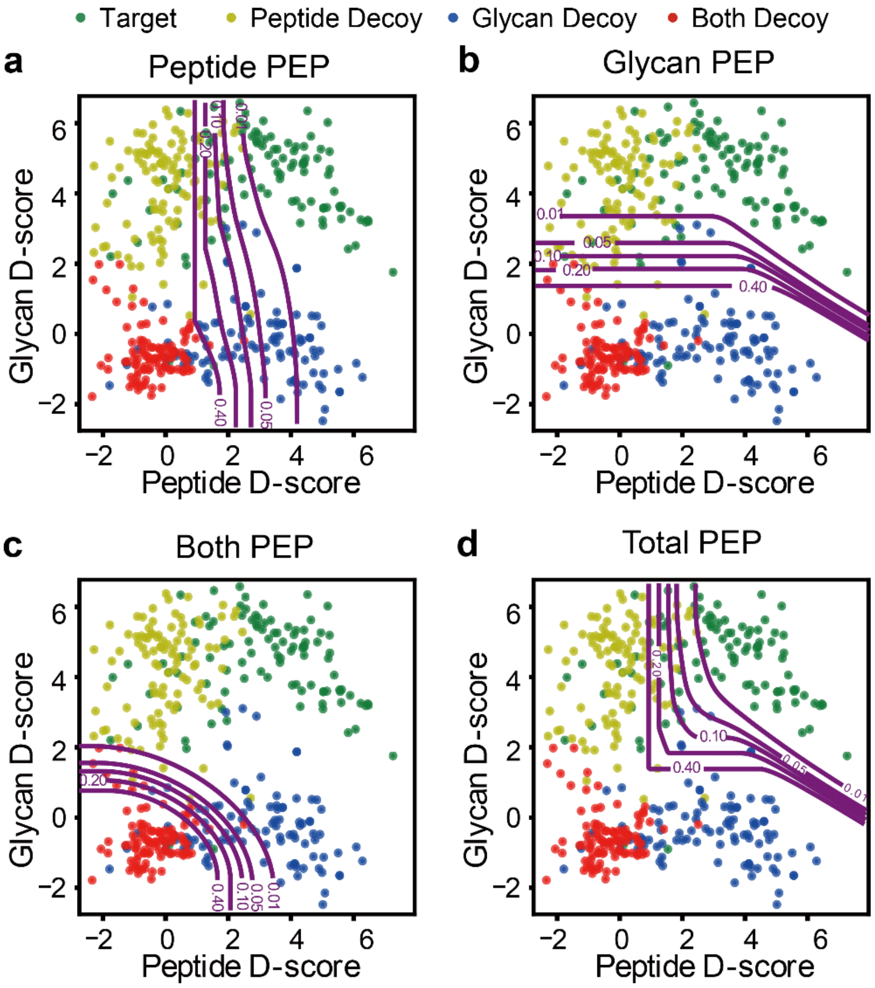


**Supplementary Fig. 2.** Contour plots of posterior error probability (PEP) of peak groups extracted from the fission yeast DIA data using the SSL library. (**a**) PEP that the peptide part of a peak group is a false identification. (**b**) PEP that the glycan part of a peak group is a false identification. (**c**) PEP that both the peptide and the glycan parts of a peak group are false identifications. (**d**) PEP that both/either the peptide and/or glycan parts of a peak group are false identifications. Green color indicates target peak groups, yellow indicates peptide decoy peak groups, blue indicates glycan decoy peak groups, and red indicates both decoy peak groups.

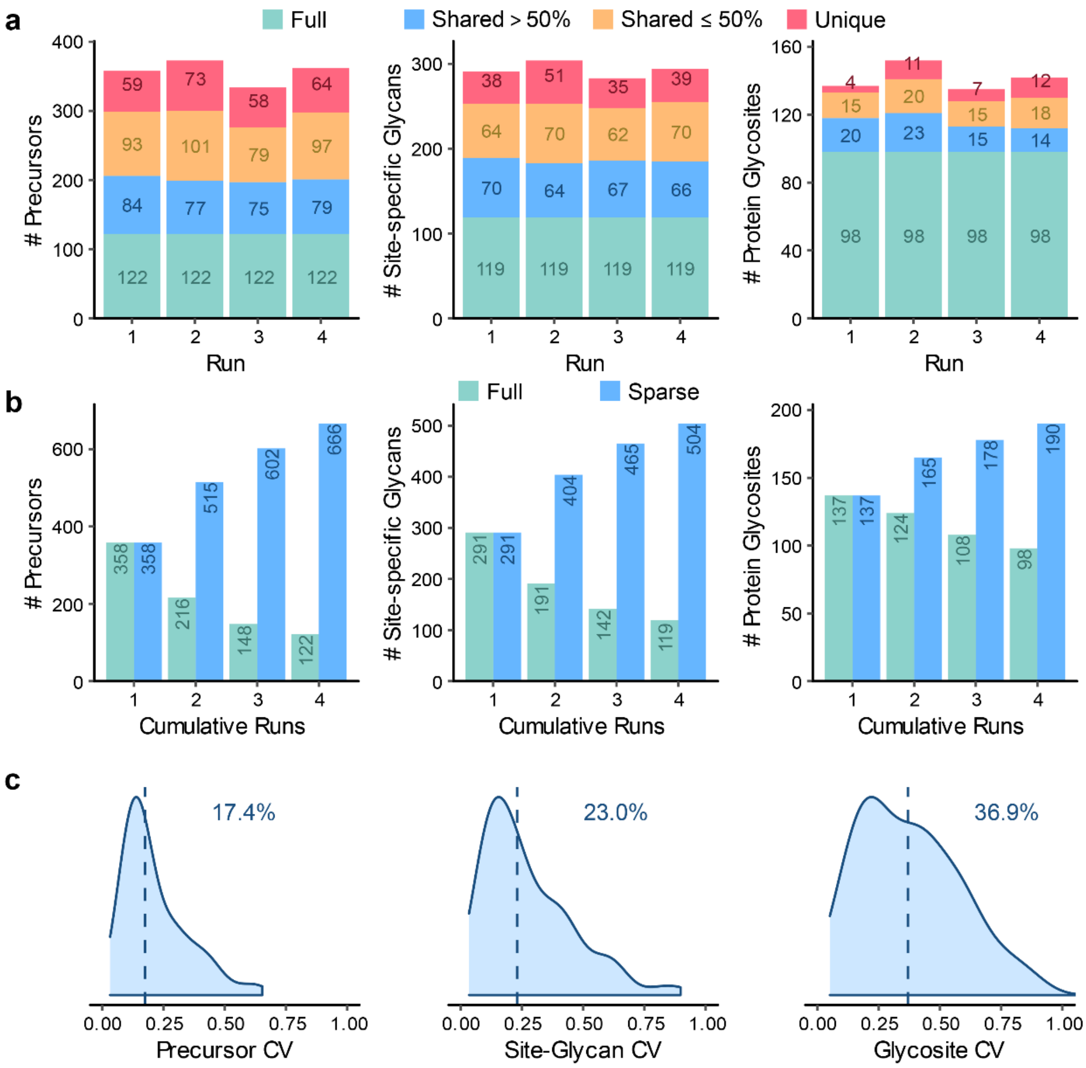


**Supplementary Fig. 3.** DDA results of the fission yeast sample with an 1 h LC gradient at the level of precursor, site-specific glycan and protein glycosite. (**a**) Numbers of identifications per run. "Full" represents identifications observed in all the runs; "shared >50%" represents identifications observed in 3 runs; "shared ≤50%" represents identifications observed in 2 runs; "unique" represents identifications observed in only 1 run. (**b**) Numbers of cumulative identifications from run 1 to 4. "Full" represents identifications shared in the cumulative runs; "sparse" represents identifications observed in at least one run in the cumulative runs. (**c**) Coefficients of variation (CVs) of quantification results. Medians are indicated.

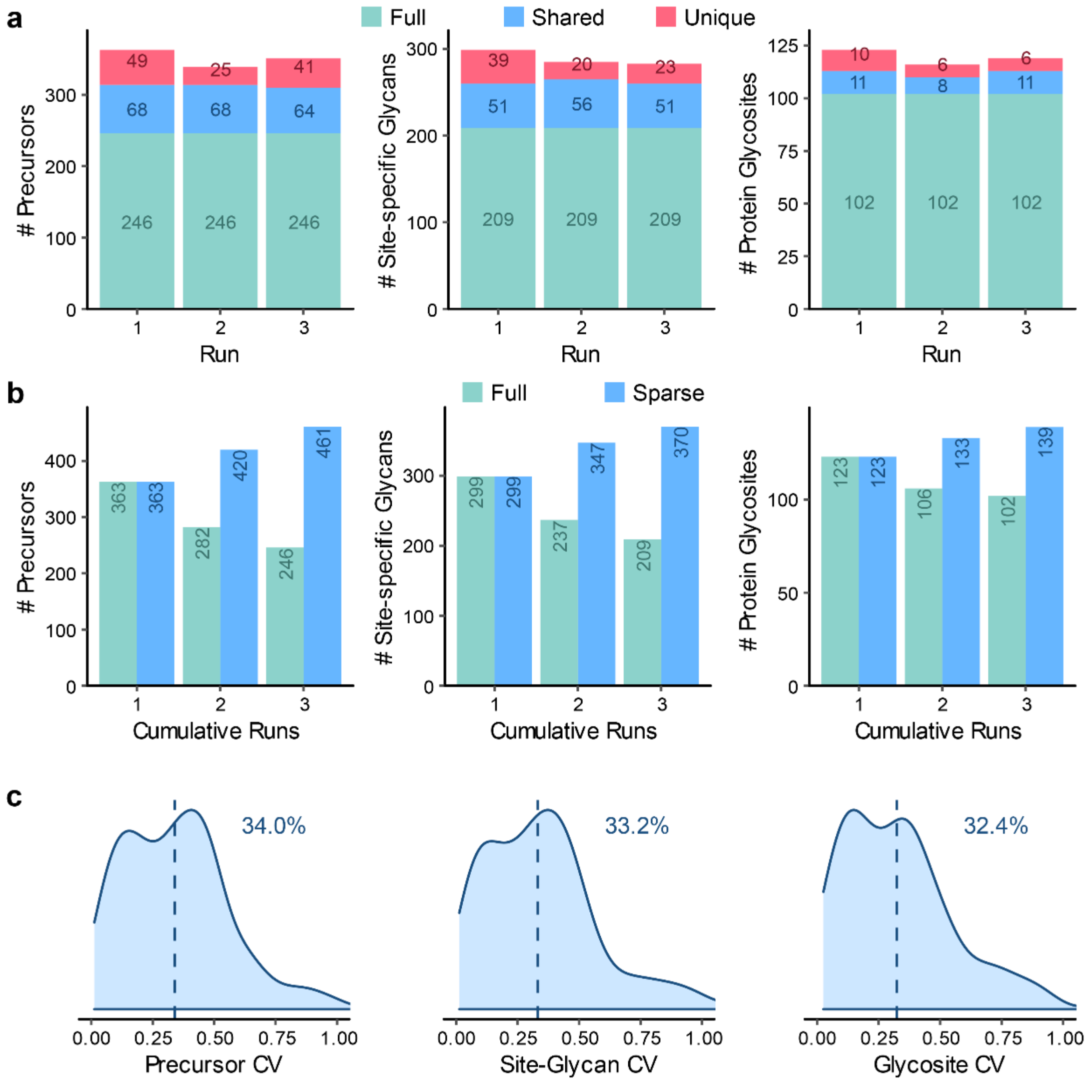


**Supplementary Fig. 4.** DDA results of the fission yeast sample with a 6 h LC gradient at the level of precursor, site-specific glycan and protein glycosite. (**a**) Numbers of identifications per run. "Full" represents identifications observed in all the runs; "shared" represents identifications observed in 2 runs; "unique" represents identifications observed in only 1 run. (**b**) Numbers of cumulative identifications from run 1 to 3. "Full" represents identifications shared in the cumulative runs; "sparse" represents identifications observed in at least one run in the cumulative runs. (**c**) Coefficients of variation (CVs) of quantification results. Medians are indicated.

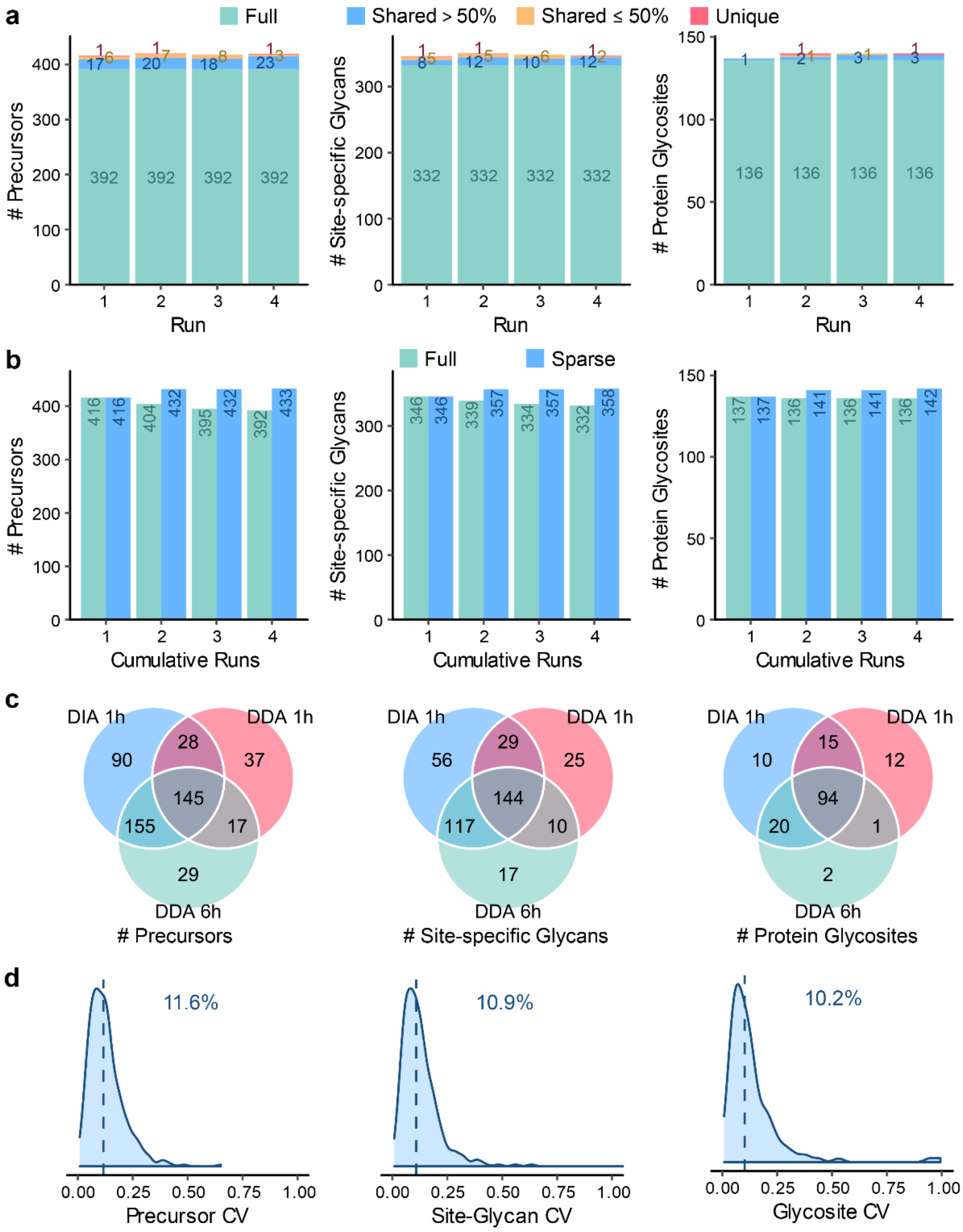


**Supplementary Fig. 5.** DIA results of the fission yeast sample using the sample-specific library at the level of precursor, site-specific glycan and protein glycosite. (**a**) Numbers of identifications per run. “Full” represents identifications observed in all the runs; “shared >50%” represents identifications observed in 3 runs; “shared ≤50%” represents identifications observed in 2 runs; “unique” represents identifications observed in only 1 run. (**b**) Numbers of cumulative identifications from run 1 to 4. “Full” represents identifications shared in the cumulative runs; “sparse” represents identifications observed in at least one run in the cumulative runs. (**c**) Comparison of numbers of identifications shared in >50% runs using DDA with an 1 h LC gradient, DDA with a 6 h LC gradient, and DIA with an 1 h LC gradient. (**d**) Coefficients of variation (CVs) of quantification results by DIA. Medians are indicated.

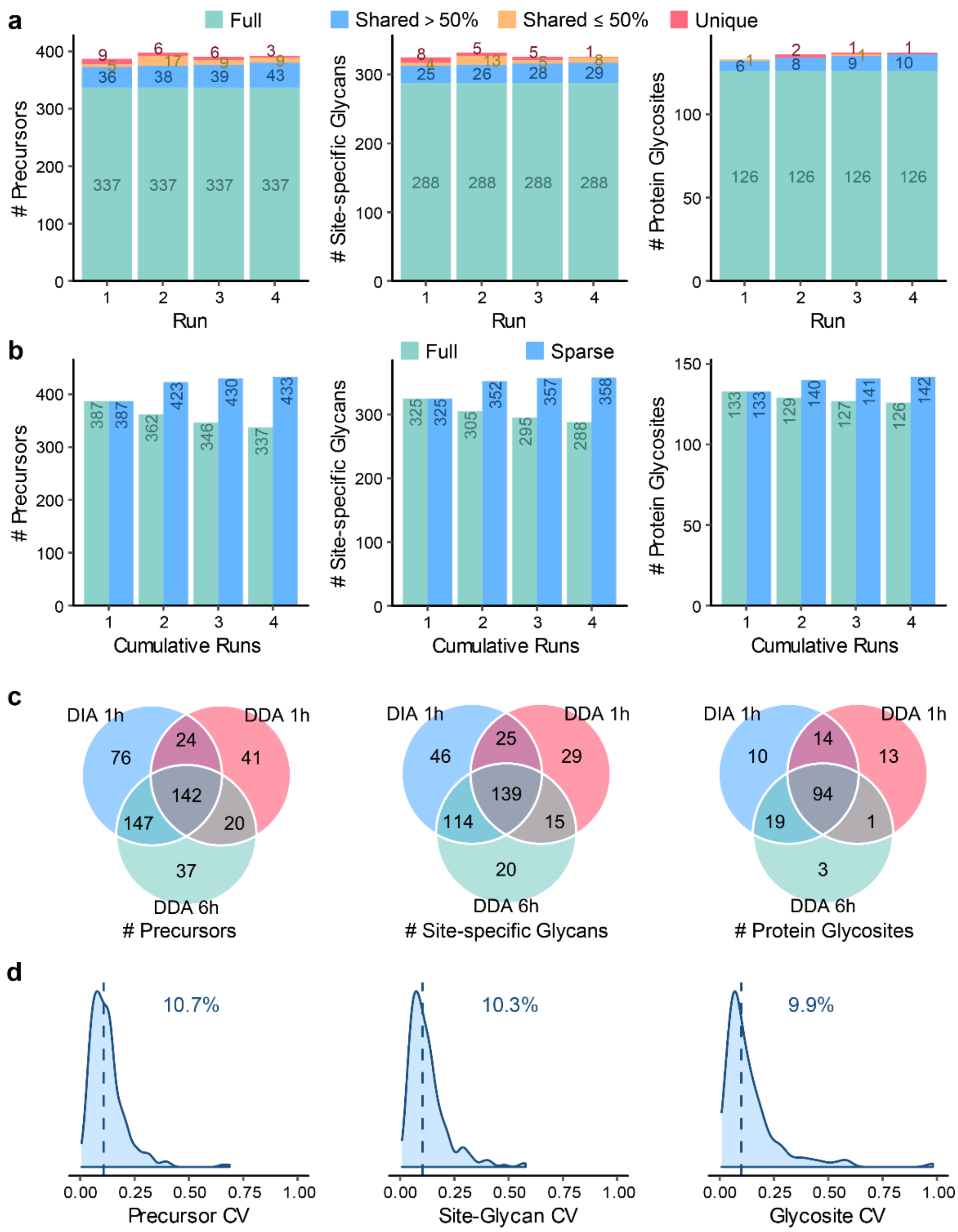


**Supplementary Fig. 6.** DIA results without multi-run alignment of the fission yeast sample using the sample-specific library at the level of precursor, site-specific glycan and protein glycosite. (**a**) Numbers of identifications per run. “Full” represents identifications observed in all the runs; “shared >50%” represents identifications observed in 3 runs; “shared ≤50%” represents identifications observed in 2 runs; “unique” represents identifications observed in only 1 run. (**b**) Numbers of cumulative identifications from run 1 to 4. “Full” represents identifications shared in the cumulative runs; “sparse” represents identifications observed in at least one run in the cumulative runs. (**c**) Comparison of numbers of identifications shared in >50% runs using DDA with an 1 h LC gradient, DDA with a 6 h LC gradient, and DIA with an 1 h LC gradient. (**d**) Coefficients of variation (CVs) of quantification results by DIA. Medians are indicated.

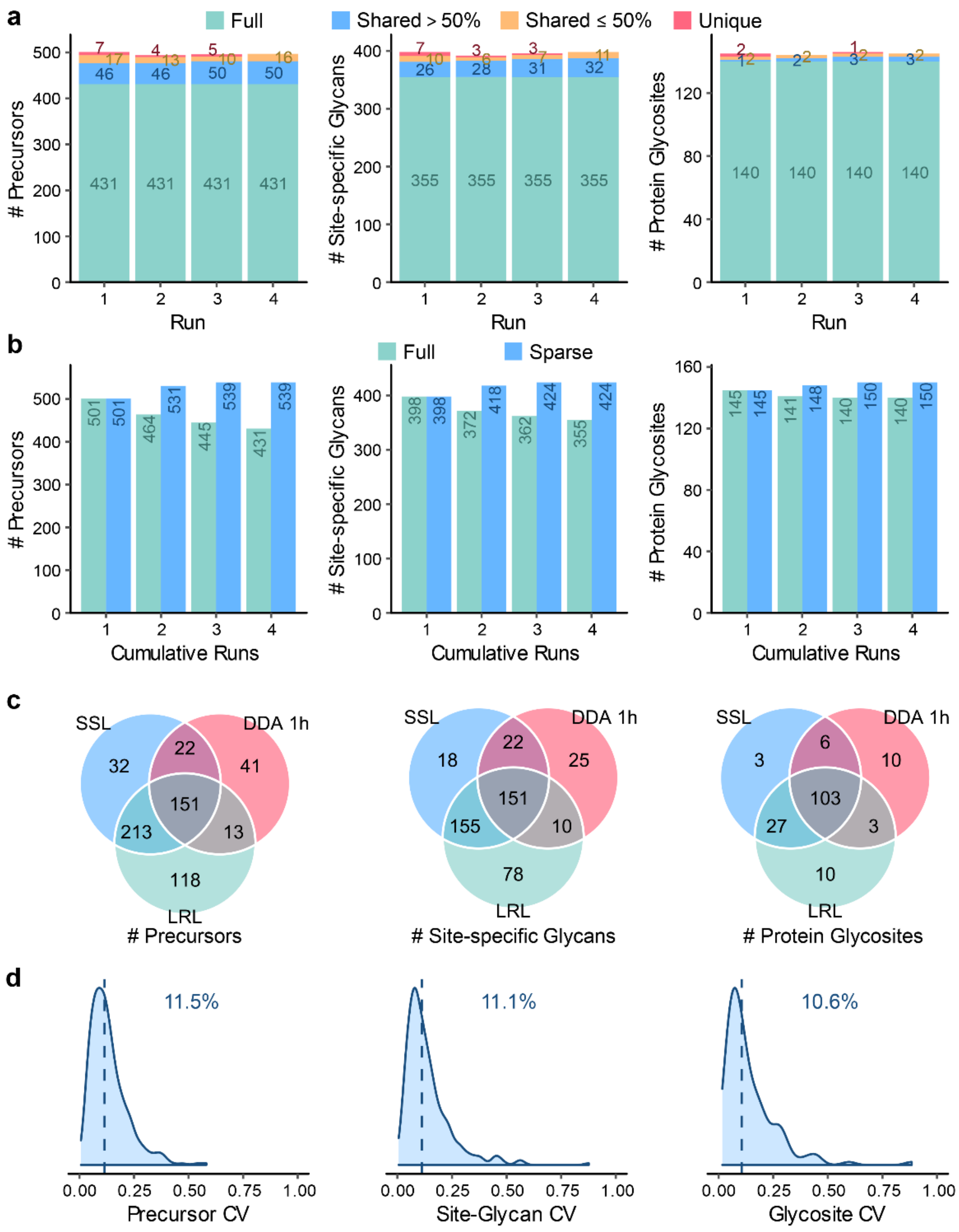


**Supplementary Fig. 7.** DIA results of the fission yeast sample using the lab repository-scale library at the level of precursor, site-specific glycan and protein glycosite. (**a**) Numbers of identifications per run. "Full" represents identifications observed in all the runs; "shared >50%" represents identifications observed in 3 runs; "shared ≤50%" represents identifications observed in 2 runs; "unique" represents identifications observed in only 1 run. (**b**) Numbers of cumulative identifications from run 1 to 4. "Full" represents identifications shared in the cumulative runs; "sparse" represents identifications observed in at least one run in the cumulative runs. (**c**) Comparison of numbers of identifications shared in >50% runs using DDA with an 1 h LC gradient, DIA with the sample-specific library (SSL), and DIA with the lab repository-scale library (LRL). (**d**) Coefficients of variation (CVs) of quantification results by DIA with LRL. Medians are indicated.

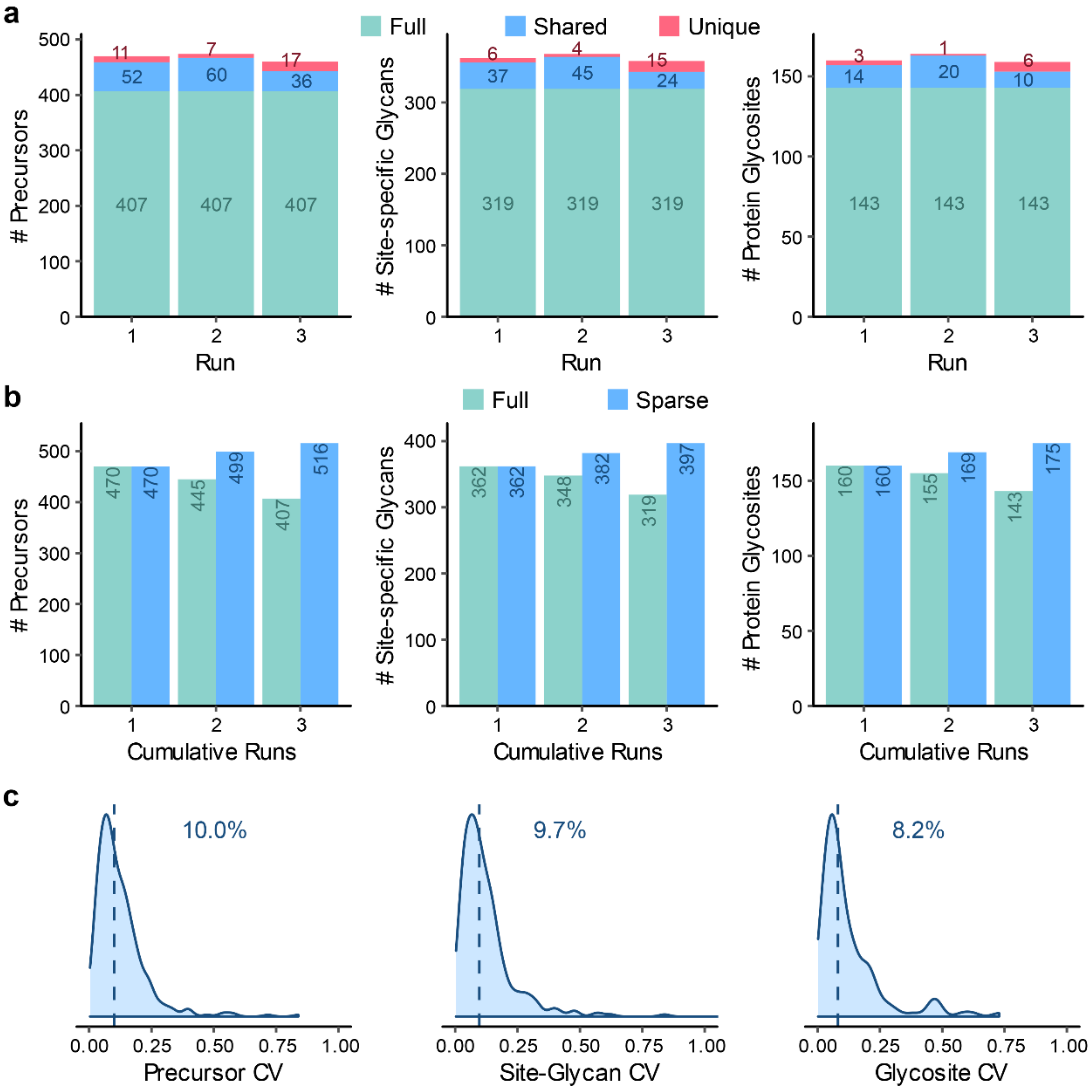


**Supplementary Fig. 8.** DIA results of the budding yeast sample at the level of precursor, site-specific glycan and protein glycosite. (**a**) Numbers of identifications per run. "Full" represents identifications observed in all the runs; "shared" represents identifications observed in 2 runs; "unique" represents identifications observed in only 1 run. (**b**) Numbers of cumulative identifications from run 1 to 3. "Full" represents identifications shared in the cumulative runs; "sparse" represents identifications observed in at least one run in the cumulative runs. (**c**) Coefficients of variation (CVs) of quantification results. Medians are indicated.

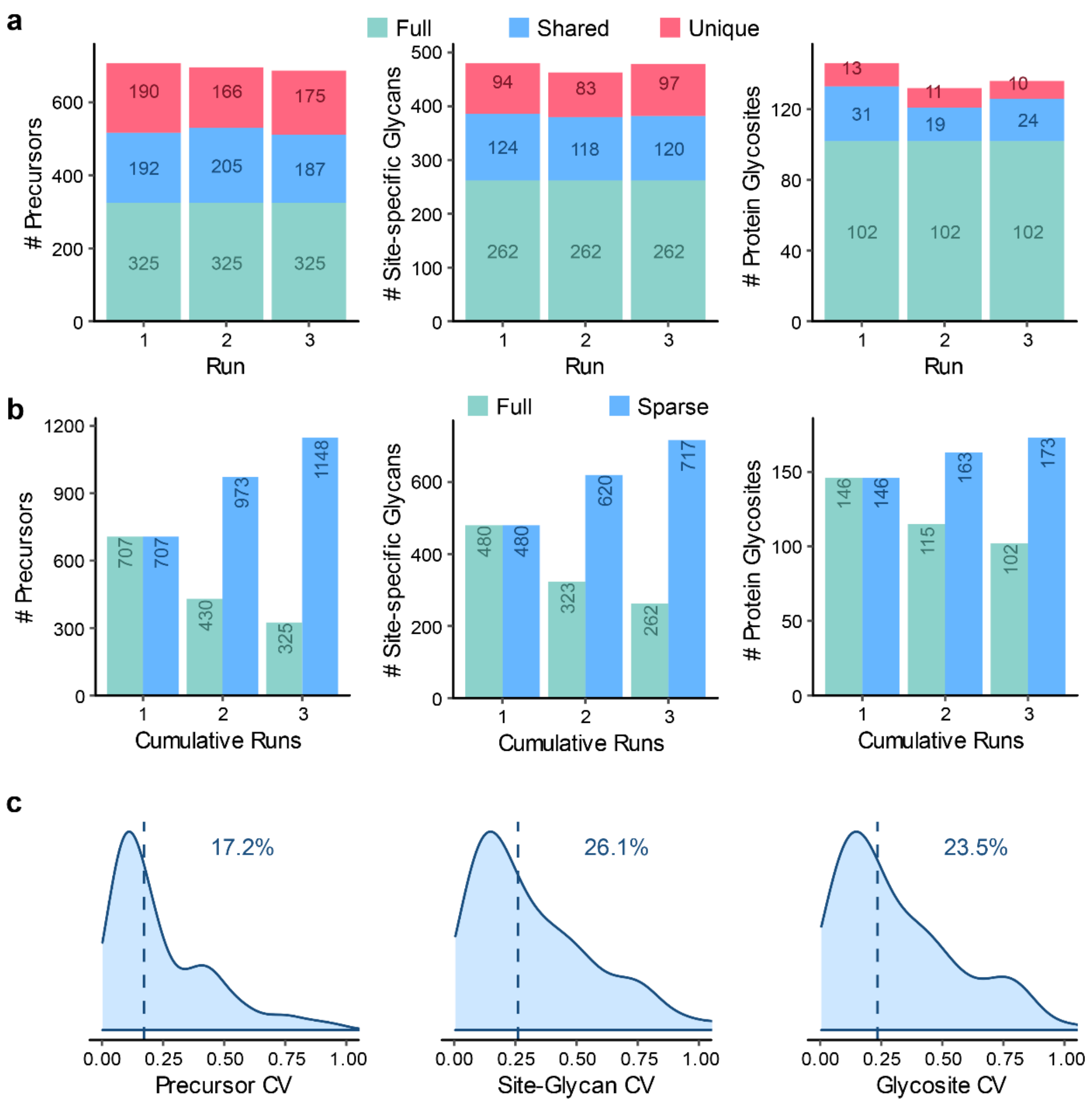


**Supplementary Fig. 9.** DDA results of the human serum sample at the level of precursor, site-specific glycan and protein glycosite. (**a**) Numbers of identifications per run. "Full" represents identifications observed in all the runs; "shared" represents identifications observed in 2 runs; "unique" represents identifications observed in only 1 run. (**b**) Numbers of cumulative identifications from run 1 to 3. "Full" represents identifications shared in the cumulative runs; "sparse" represents identifications observed in at least one run in the cumulative runs. (**c**) Coefficients of variation (CVs) of quantification results. Medians are indicated.

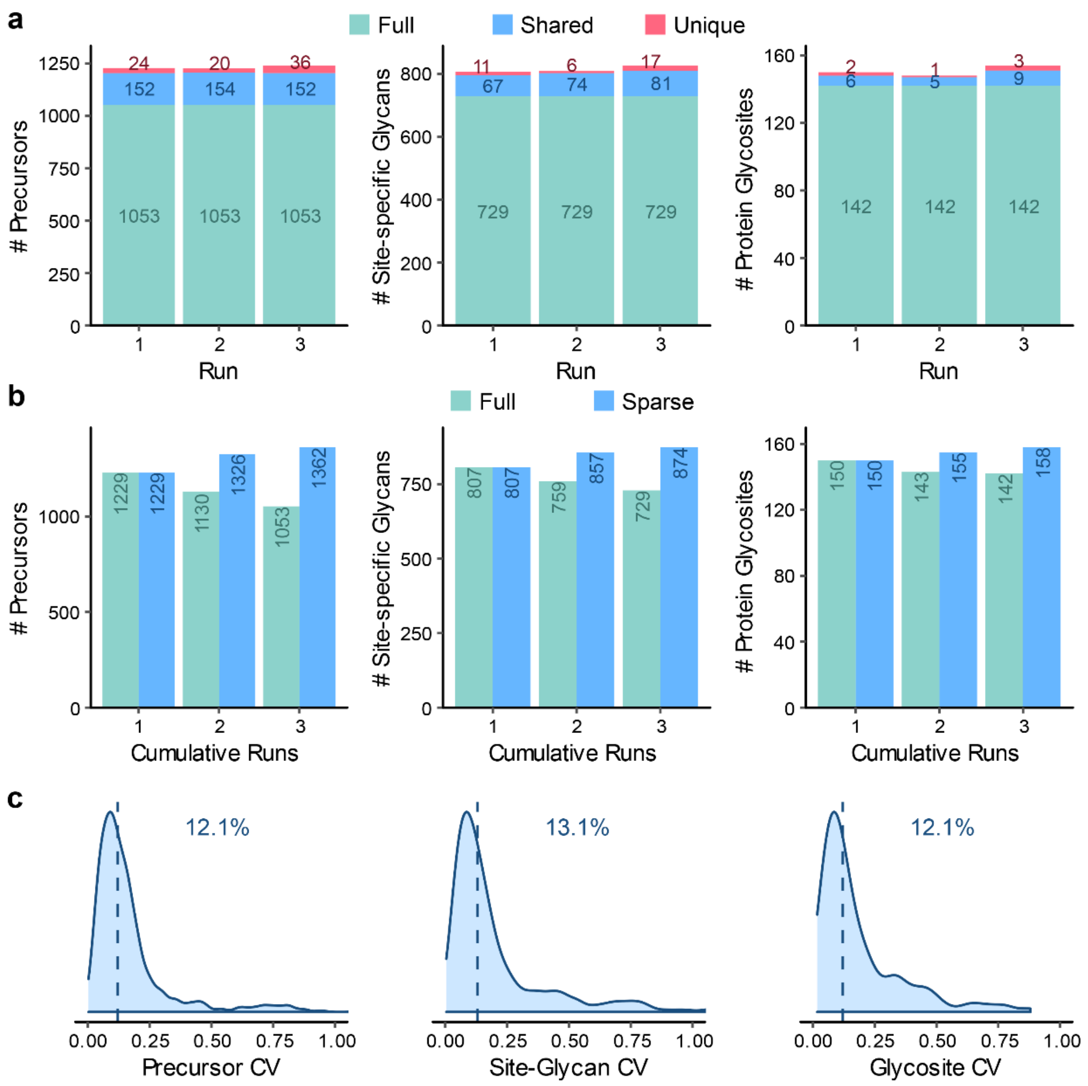


**Supplementary Fig. 10.** DIA results without glycoform inference of the human serum sample using the sample-specific library at the level of precursor, site-specific glycan and protein glycosite. (**a**) Numbers of identifications per run. "Full" represents identifications observed in all the runs; "shared" represents identifications observed in 2 runs; "unique" represents identifications observed in only 1 run. (**b**) Numbers of cumulative identifications from run 1 to 3. "Full" represents identifications shared in the cumulative runs; "sparse" represents identifications observed in at least one run in the cumulative runs. (**c**) Coefficients of variation (CVs) of quantification results. Medians are indicated.

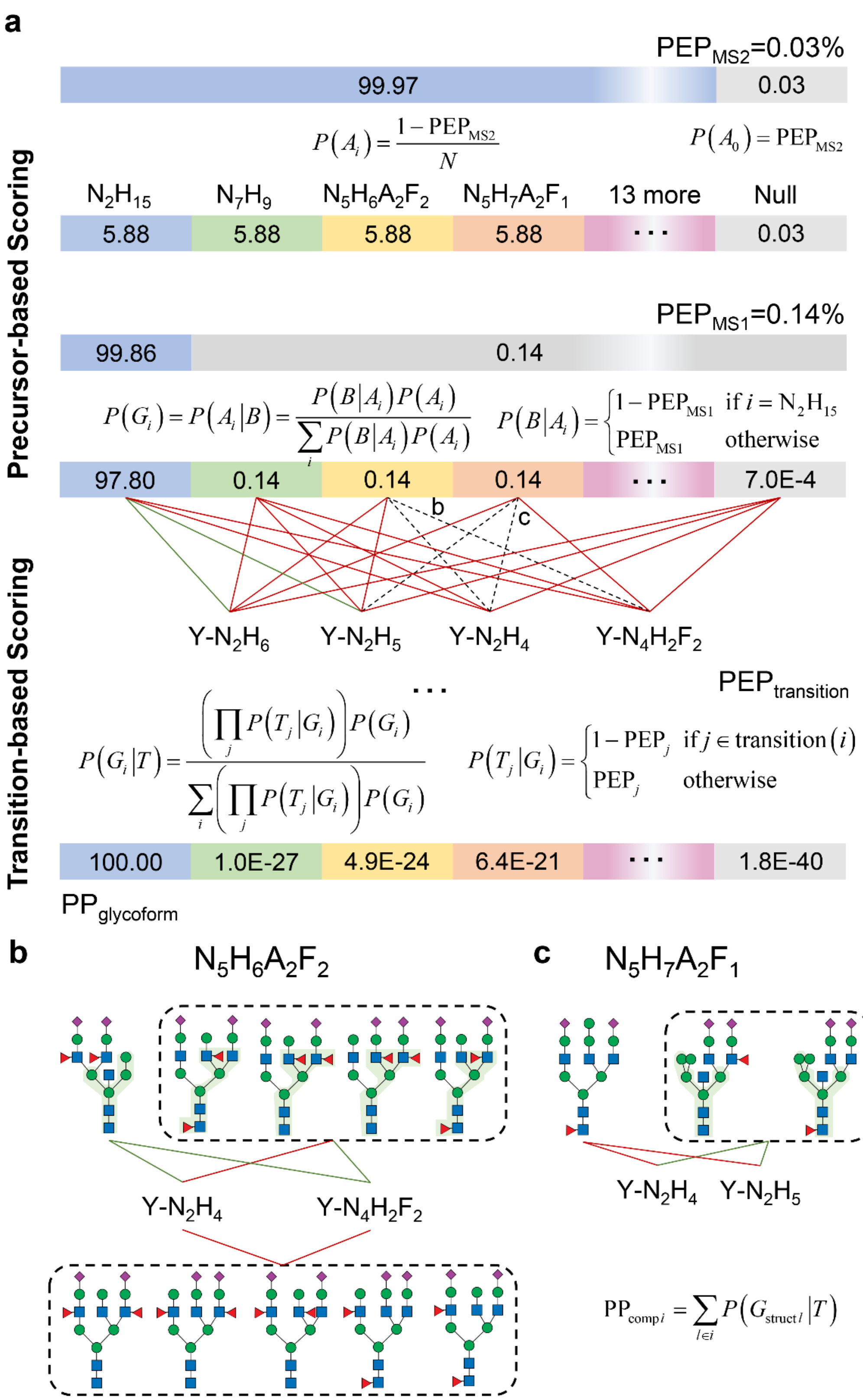


**Supplementary Fig. 11.** The Bayesian hierarchical model for glycoform inference. (**a**) Integrating the precursor and transition posterior probabilities (PPs) to calculate glycoform PP. (**b**) Isomeric glycan structures of $HexNAc_5Hex_6NeuAc_2Fuc_2$. (**c**) Isomeric glycan structures of $HexNAc_5Hex_7NeuAc_2Fuc_1$. The glycan symbols are as follows: a green circle or "H" represents Hex; a blue square or "N" represents HexNAc;

a red triangle or “F” represents Fuc; a purple diamond or “A” represents NeuAc. A green line indicates a Y ion can be originated from a glycoform (glycan composition or structure); otherwise, they are connected with a red line. The information for the dashed lines is expanded in (**b**) and (**c**) because different isomeric glycan structures of the corresponding glycoforms can generate different fragment ions. PEP: posterior error probabilities.

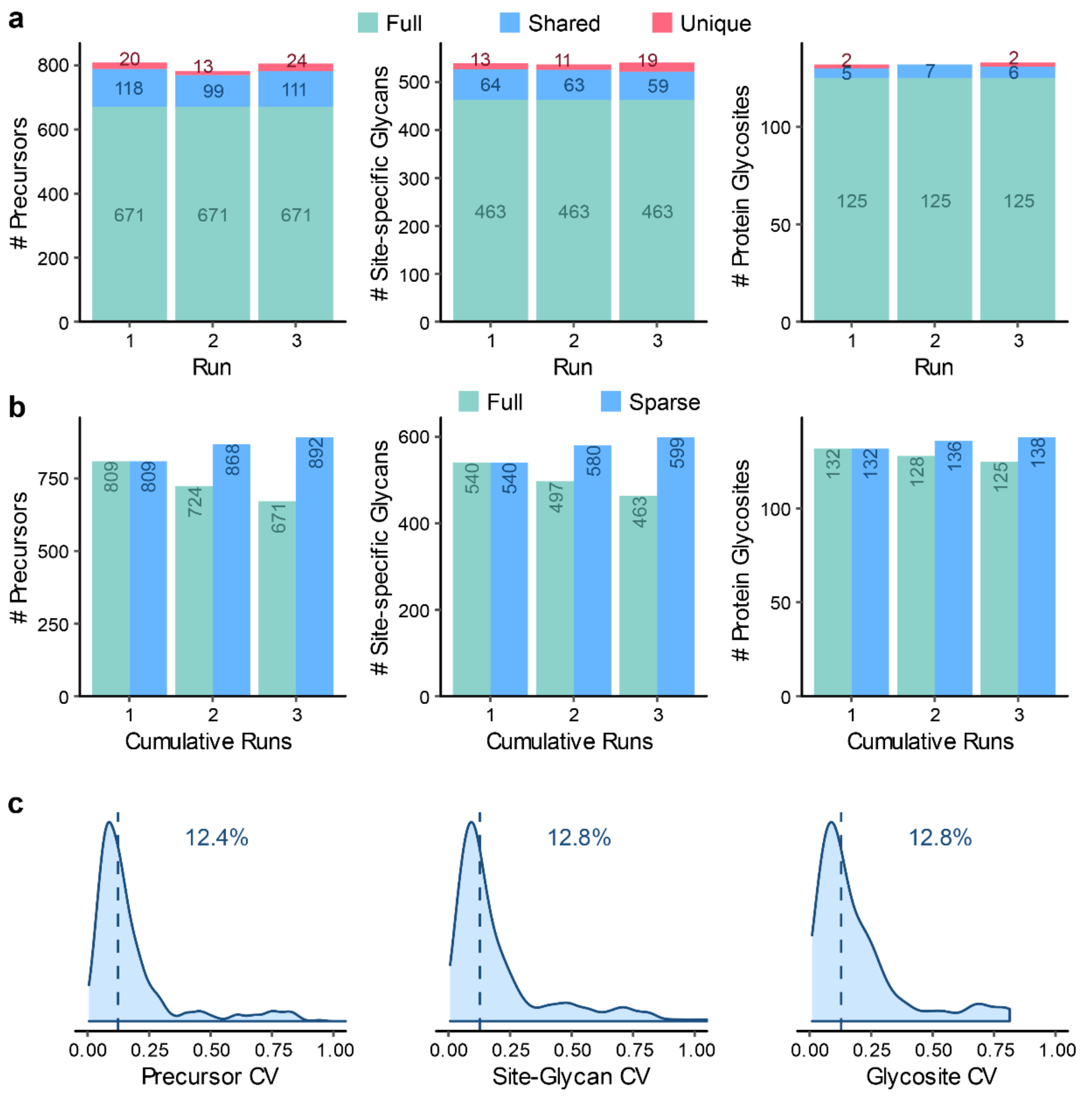


**Supplementary Fig. 12.** DIA results with glycoform inference of the human serum sample using the sample-specific library at the level of precursor, site-specific glycan and protein glycosite. (**a**) Numbers of identifications per run. “Full” represents identifications observed in all the runs; “shared” represents identifications observed in 2 runs; “unique” represents identifications observed in only 1 run. (**b**) Numbers of cumulative identifications from run 1 to 3. “Full” represents identifications shared in the cumulative runs; “sparse” represents identifications observed in at least one run in the cumulative runs. (**c**) Coefficients of variation (CVs) of quantification results. Medians are indicated.

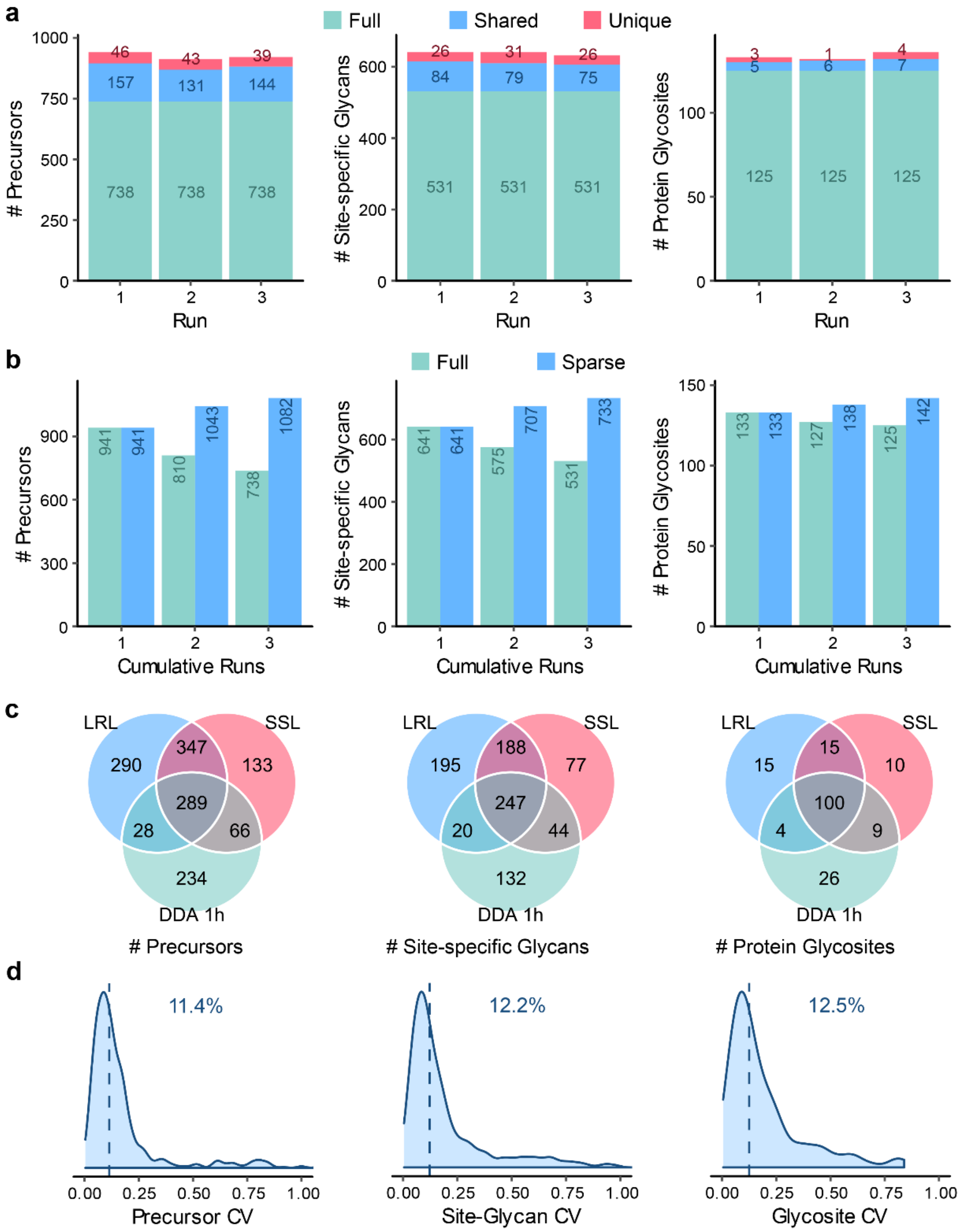


**Supplementary Fig. 13.** DIA results with glycoform inference of the human serum sample using the lab repository-scale library at the level of precursor, site-specific glycan and protein glycosite. (**a**) Numbers of identifications per run. "Full" represents identifications observed in all the runs; "shared" represents identifications observed in 2 runs; "unique" represents identifications observed in only 1 run. (**b**) Numbers of cumulative identifications from run 1 to 3. "Full" represents identifications shared in the cumulative runs; "sparse" represents identifications observed in at least one run in the cumulative runs. (**c**) Comparison of numbers of identifications shared in >50% runs using DDA, DIA with the sample-specific library (SSL), and DIA with the lab repository-scale library (LRL). (**d**) Coefficients of variation (CVs) of quantification results by DIA with LRL. Medians are indicated.

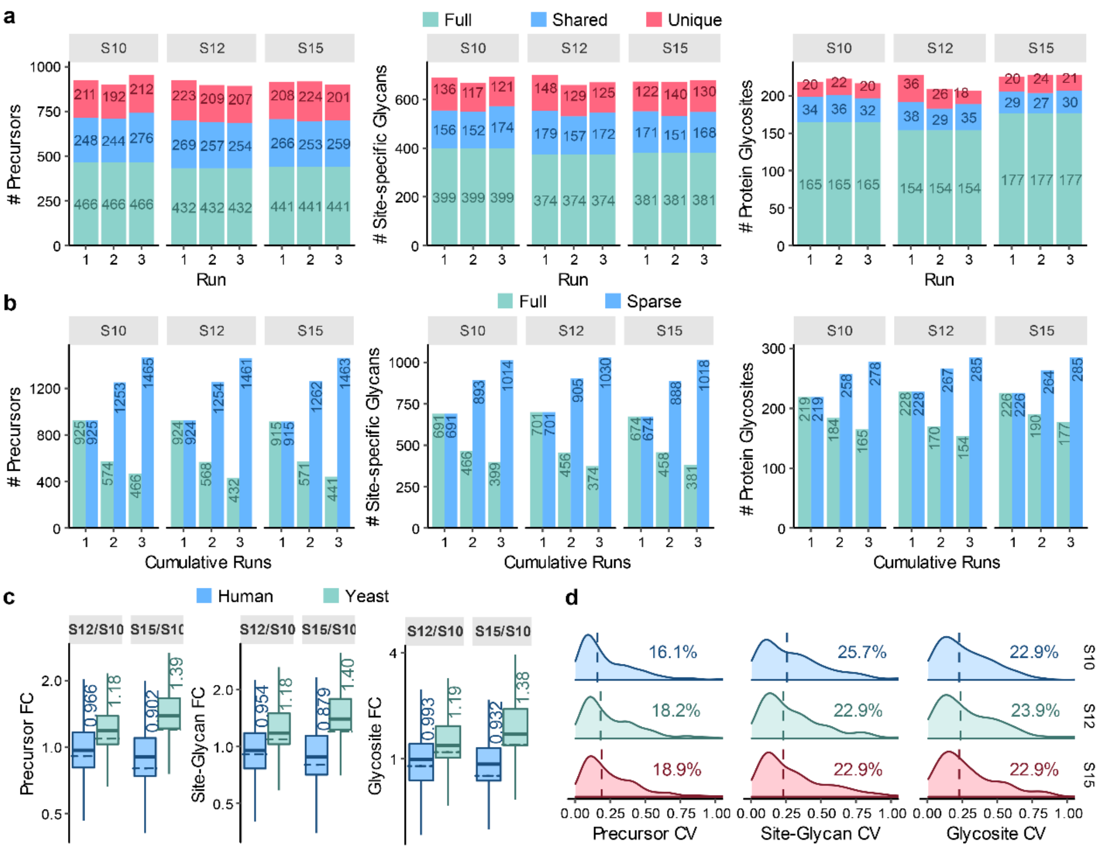


**Supplementary Fig. 14.** DDA results of the mixed-organism samples at the level of precursor, site-specific glycan and protein glycosite. (**a**) Numbers of identifications per run of each sample. "Full" represents identifications observed in all the runs; "shared" represents identifications observed in 2 runs; "unique" represents identifications observed in only 1 run. (**b**) Numbers of cumulative identifications from run 1 to 3 of each sample. "Full" represents identifications shared in the cumulative runs; "sparse" represents identifications observed in at least one run in the cumulative runs. (**c**) Box plot visualization of fold change of the quantification results of the mixed-organism samples. Percent changes were calculated based on the mean quantities in three replicates of each sample. The medians are indicated. The boxes indicate the interquartile ranges (IQR), and whiskers indicate 1.5 × IQR values; no outliers are shown. The dashed lines indicate theoretical fold changes of the organisms (1:0.9:0.8 (S10:S12:S15) for human and 1:1.1:1.2 (S10:S12:S15) for yeast). (**d**) Coefficients of variation (CVs) of quantification results of each sample. Medians are indicated.

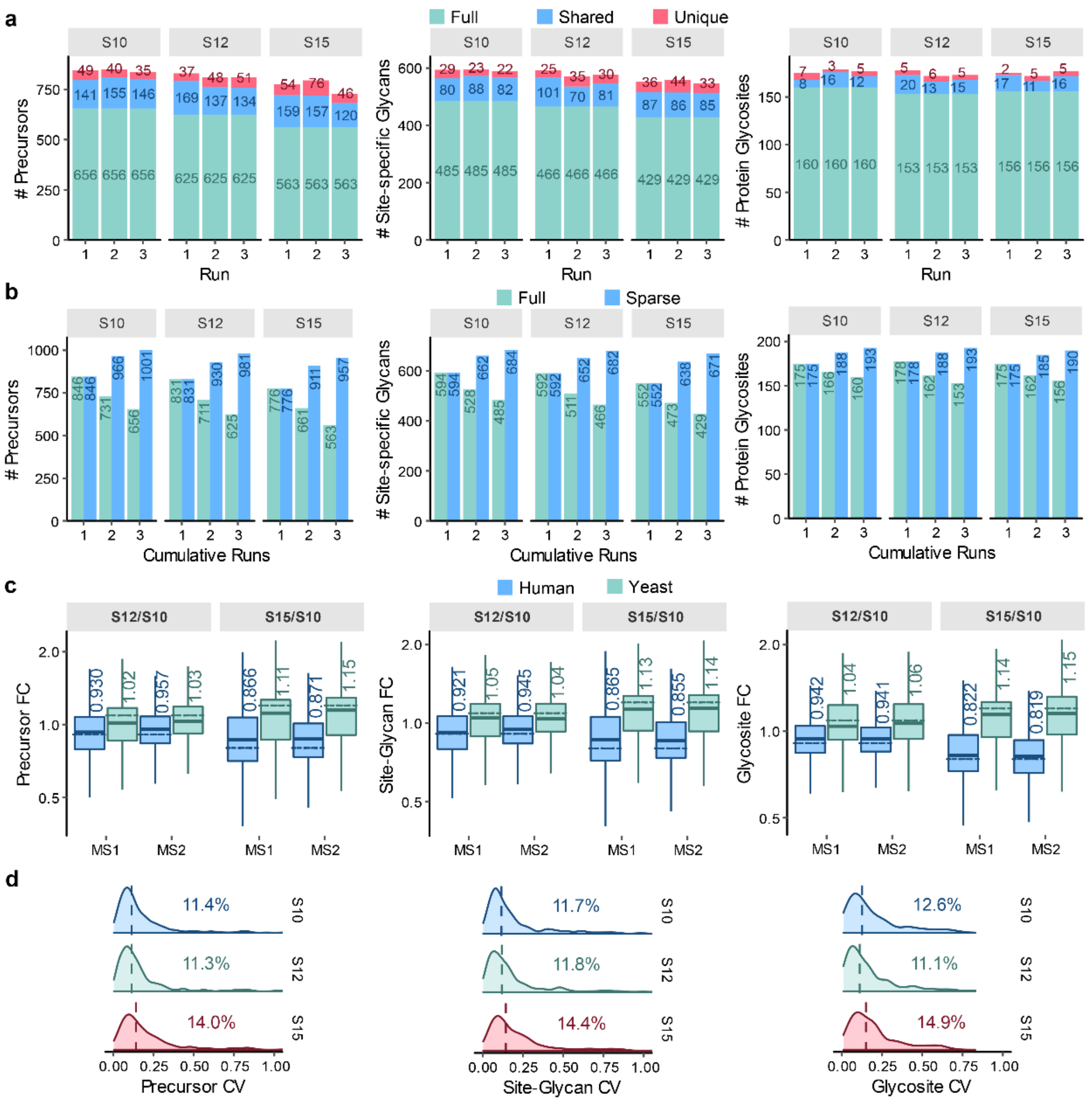


**Supplementary Fig. 15.** DIA results of the mixed-organism samples using a combined library of the budding yeast library and the serum sample-specific library (SSL) at the level of precursor, site-specific glycan and protein glycosite. (**a**) Numbers of identifications per run of each sample. “Full” represents identifications observed in all the runs; “shared” represents identifications observed in 2 runs; “unique” represents identifications observed in only 1 run. (**b**) Numbers of cumulative identifications from run 1 to 3 of each sample. “Full” represents identifications shared in the cumulative runs; “sparse” represents identifications observed in at least one run in the cumulative runs. (**c**) Box plot visualization of fold change of the MS1 and MS2-level quantification results of the mixed-organism samples. Percent changes were calculated based on the mean quantities in three replicates of each sample. The medians are indicated. The boxes indicate the interquartile ranges (IQR), and whiskers indicate 1.5 × IQR values; no outliers are shown. The dashed lines indicate theoretical fold changes of the organisms (1:0.9:0.8 (S10:S12:S15) for human and 1:1.1:1.2 (S10:S12:S15) for yeast). (**d**) Coefficients of variation (CVs) of quantification results of each sample. Medians are indicated.

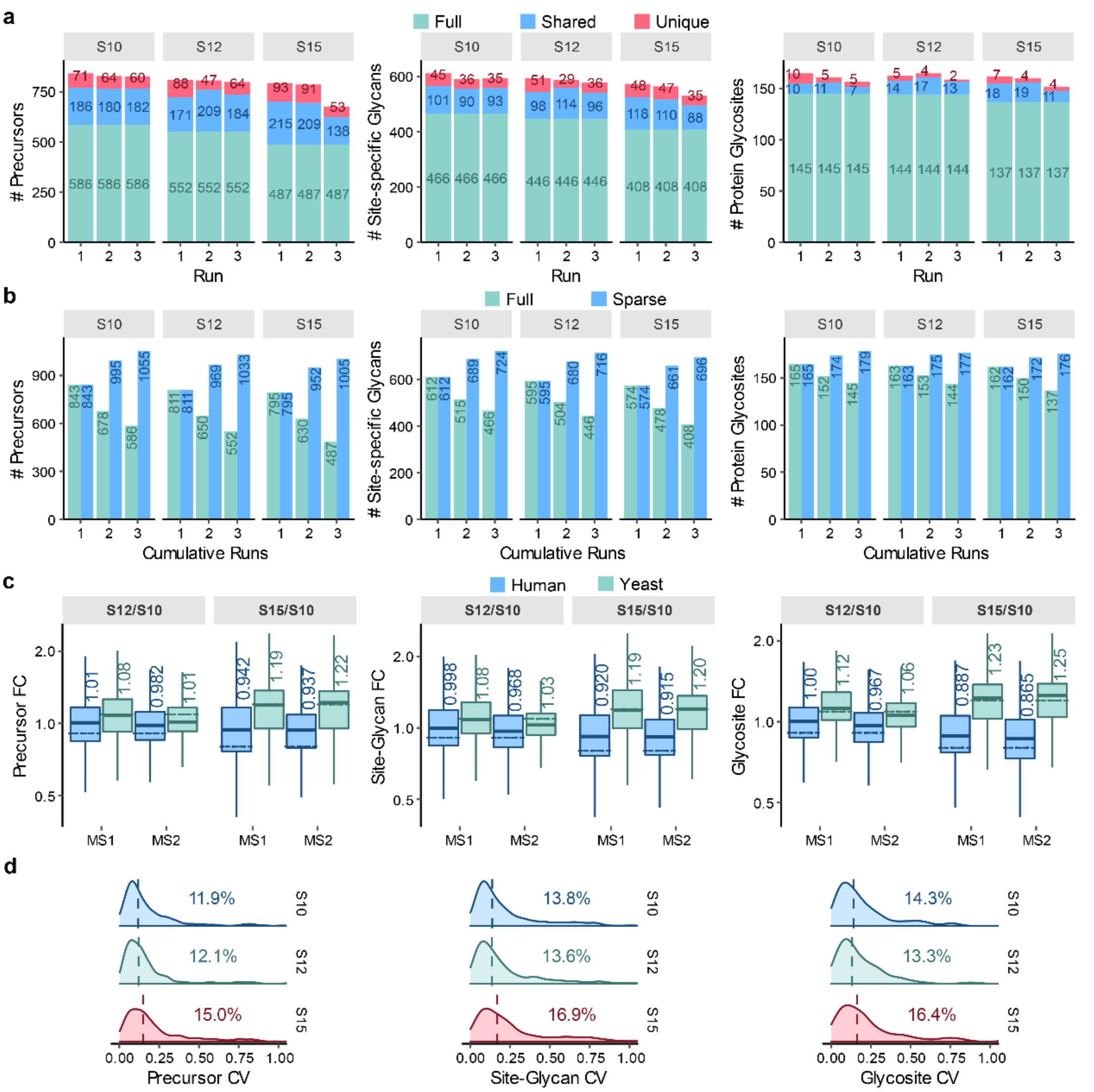


**Supplementary Fig. 16.** DIA results of the mixed-organism samples using a combined library of the budding yeast library and the serum lab repository-scale library (LRL) at the level of precursor, site-specific glycan and protein glycosite. (**a**) Numbers of identifications per run of each sample. “Full” represents identifications observed in all the runs; “shared” represents identifications observed in 2 runs; “unique” represents identifications observed in only 1 run. (**b**) Numbers of cumulative identifications from run 1 to 3 of each sample. “Full” represents identifications shared in the cumulative runs; “sparse” represents identifications observed in at least one run in the cumulative runs. (**c**) Box plot visualization of fold change of the MS1 and MS2-level quantification results of the mixed-organism samples. Percent changes were calculated based on the mean quantities in three replicates of each sample. The medians are indicated. The boxes indicate the interquartile ranges (IQR), and whiskers indicate 1.5 × IQR values; no outliers are shown. The dashed lines indicate theoretical fold changes of the organisms (1:0.9:0.8 (S10:S12:S15) for human and 1:1.1:1.2 (S10:S12:S15) for yeast). (**d**) Coefficients of variation (CVs) of quantification results of each sample. Medians are indicated.

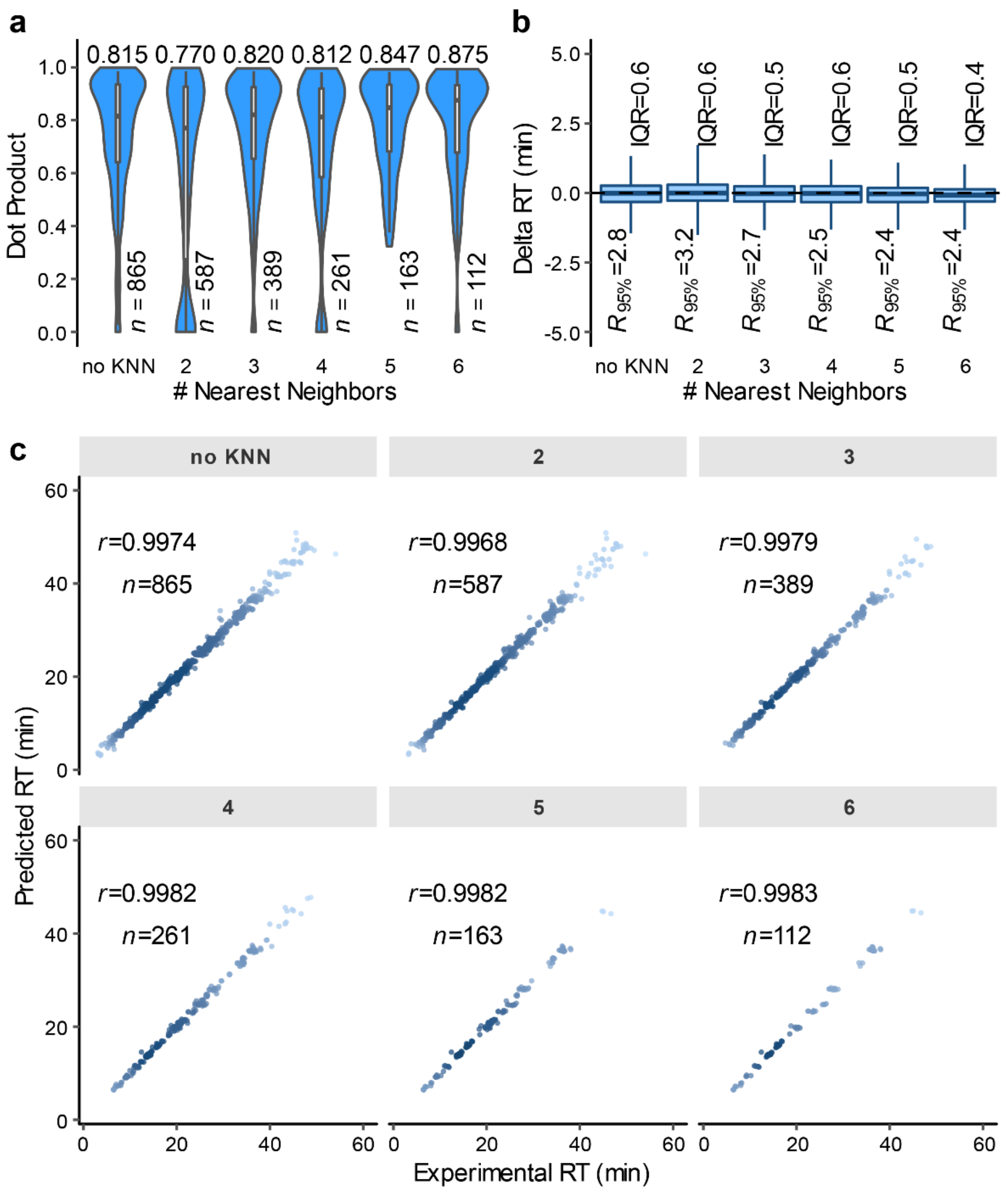


**Supplementary Fig. 17.** Cross validations of semi-empirical spectral library generation using the fission yeast lab repository-scale library (LRL), tested with different numbers of nearest neighbors *k* or without KNN. (**a**) The distributions of dot products computed between predicted and experimental MS/MS peak intensities. (**b**) The differences between predicted and experimental retention times (RTs). The medians are indicated. The boxes indicate the interquartile ranges (IQR), and the whiskers show the ranges between 2.5% and 97.5% percentiles ($R_{95\%}$); no outliers are shown. (**c**) Pearson correlation coefficients (*r*) between predicted and experimental RTs. "*n*" indicates the number of generated MS2 spectra or RT values by the prediction method.

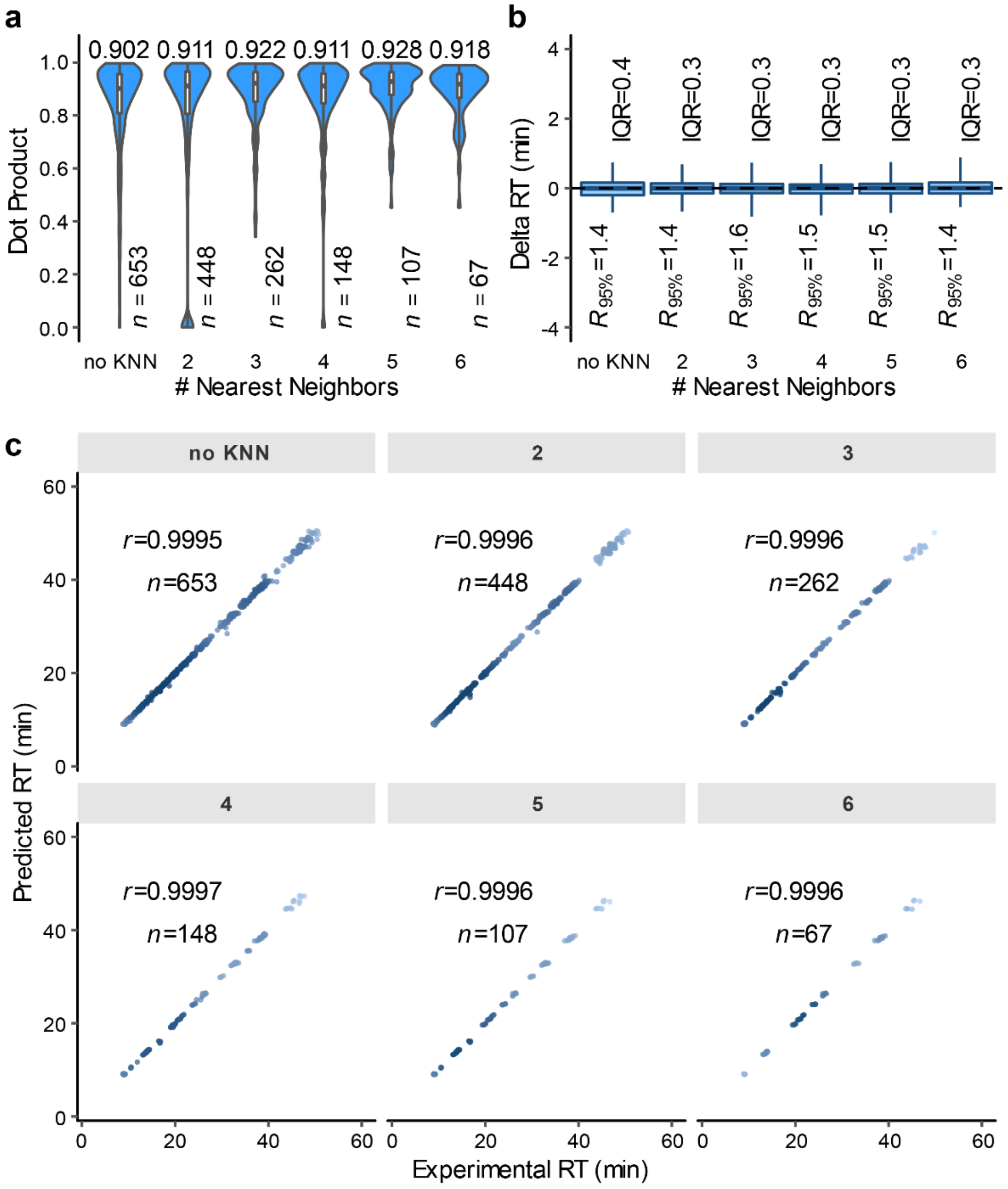


**Supplementary Fig. 18.** Cross validations of semi-empirical spectral library generation using the budding yeast library, tested with different numbers of nearest neighbors *k* or without KNN. (**a**) The distributions of dot products computed between predicted and experimental MS/MS peak intensities. (**b**) The differences between predicted and experimental retention times (RTs). The medians are indicated. The boxes indicate the interquartile ranges (IQR), and the whiskers show the ranges between 2.5% and 97.5% percentiles ($R_{95\%}$); no outliers are shown. (**c**) Pearson correlation coefficients ($r$) between predicted and experimental RTs. "*n*" indicates the number of generated MS2 spectra or RT values by the prediction method.

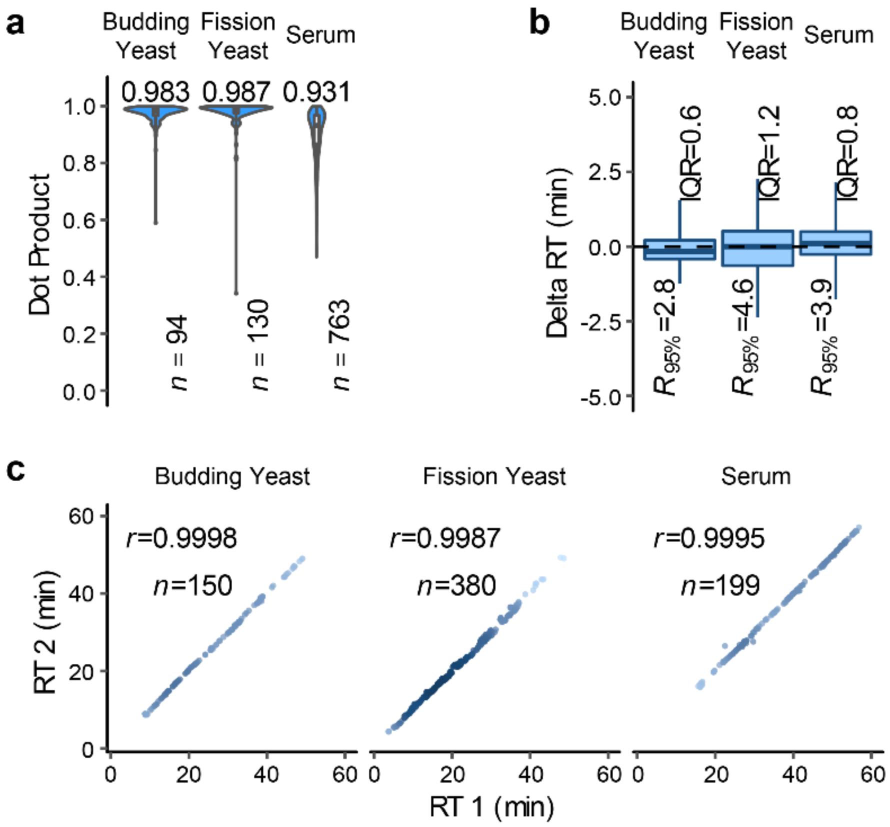


**Supplementary Fig. 19.** Statistics of replicate experimental spectra in the budding yeast, fission yeast, and serum library. Replicates within each run were combined into consensus spectra, and statistics were computed between the consensus spectra pairwise across runs. For each glycopeptide precursor, the minimum of the paired DPs and the maximum (absolute value) of the paired retention time (RT) differences were kept. (**a**) The distributions of dot products computed between MS/MS peak intensities of consensus replicate spectra. (**b**) The differences between retention times (RTs) of consensus replicate spectra. The medians are indicated. The boxes indicate the interquartile ranges (IQR), and the whiskers show the ranges between 2.5% and 97.5% percentiles ($R_{95\%}$); no outliers are shown. (**c**) Pearson correlation coefficients ($r$) between RTs of consensus replicate spectra in two randomly chosen runs.

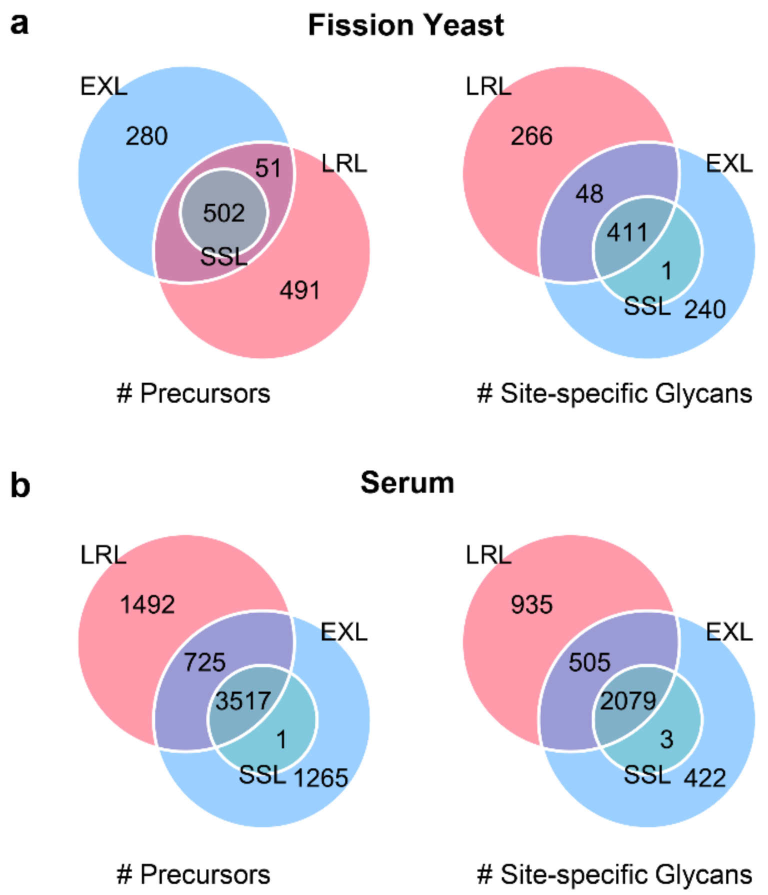


**Supplementary Fig. 20.** Coverage comparison of the sample-specific library (SSL), the lab repository-scale library (LRL), and the extended library by the semi-empirical approach (EXL). (**a**) Libraries for the fission yeast sample. (**b**) Libraries for the human serum sample.

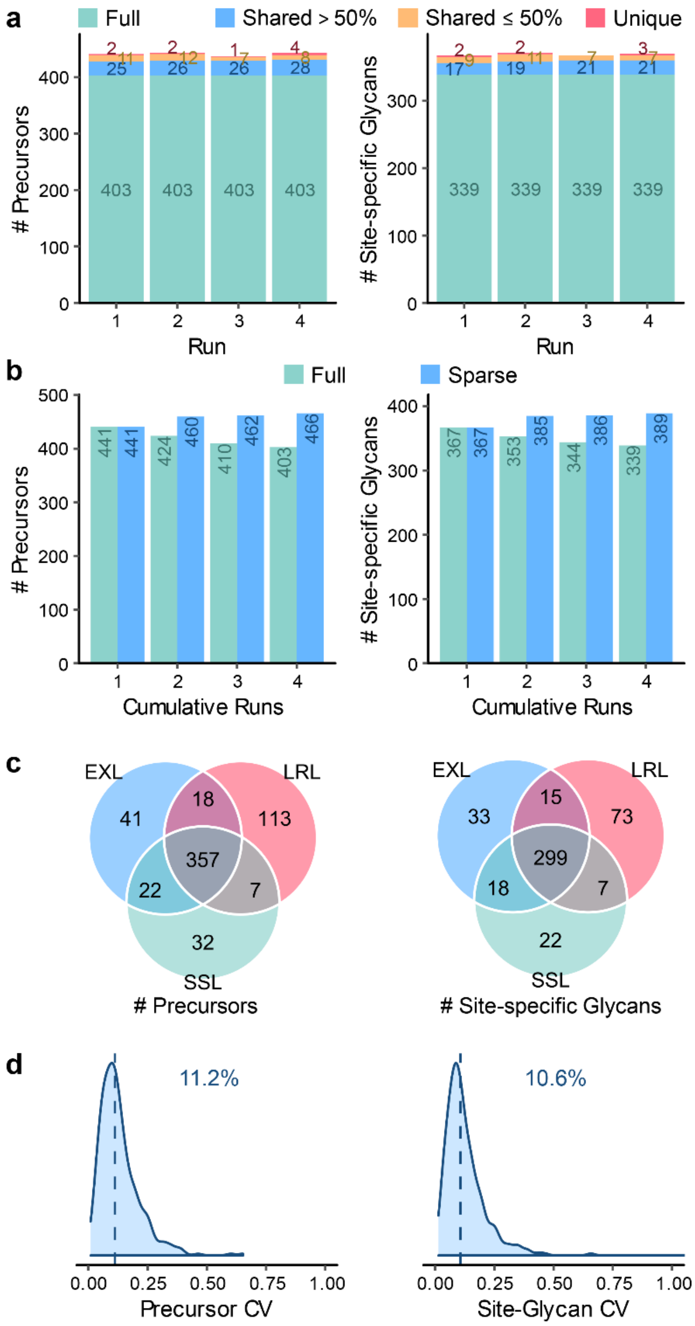


**Supplementary Fig. 21.** DIA results of the fission yeast sample using the extended library at the level of precursor and site-specific glycan. (**a**) Numbers of identifications per run. "Full" represents identifications observed in all the runs; "shared >50%" represents identifications observed in 3 runs; "shared ≤50%" represents identifications observed in 2 runs; "unique" represents identifications observed in only 1 run. (**b**) Numbers of cumulative identifications from run 1 to 4. "Full" represents identifications shared in the cumulative runs; "sparse" represents identifications observed in at least one run in the cumulative runs. (**c**) Comparison of numbers of identifications shared in >50% runs using the sample-specific library (SSL), the lab repository-scale library (LRL), and the extended library (EXL). (**d**) Coefficients of variation (CVs) of quantification results by DIA with EXL. Medians are indicated.

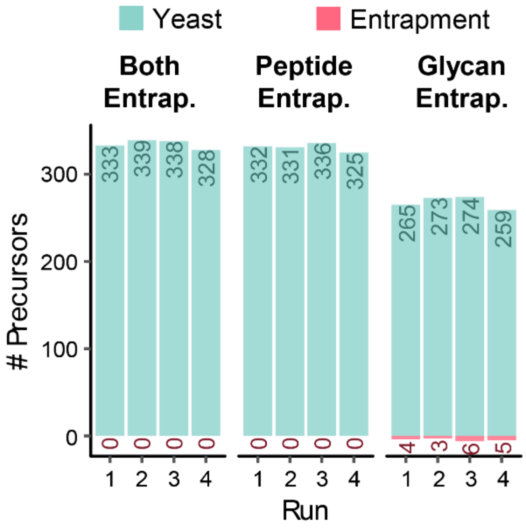


**Supplementary Fig. 22.** Numbers of identifications from the fission yeast sample using the extend libraries with entrapment glycopeptides.

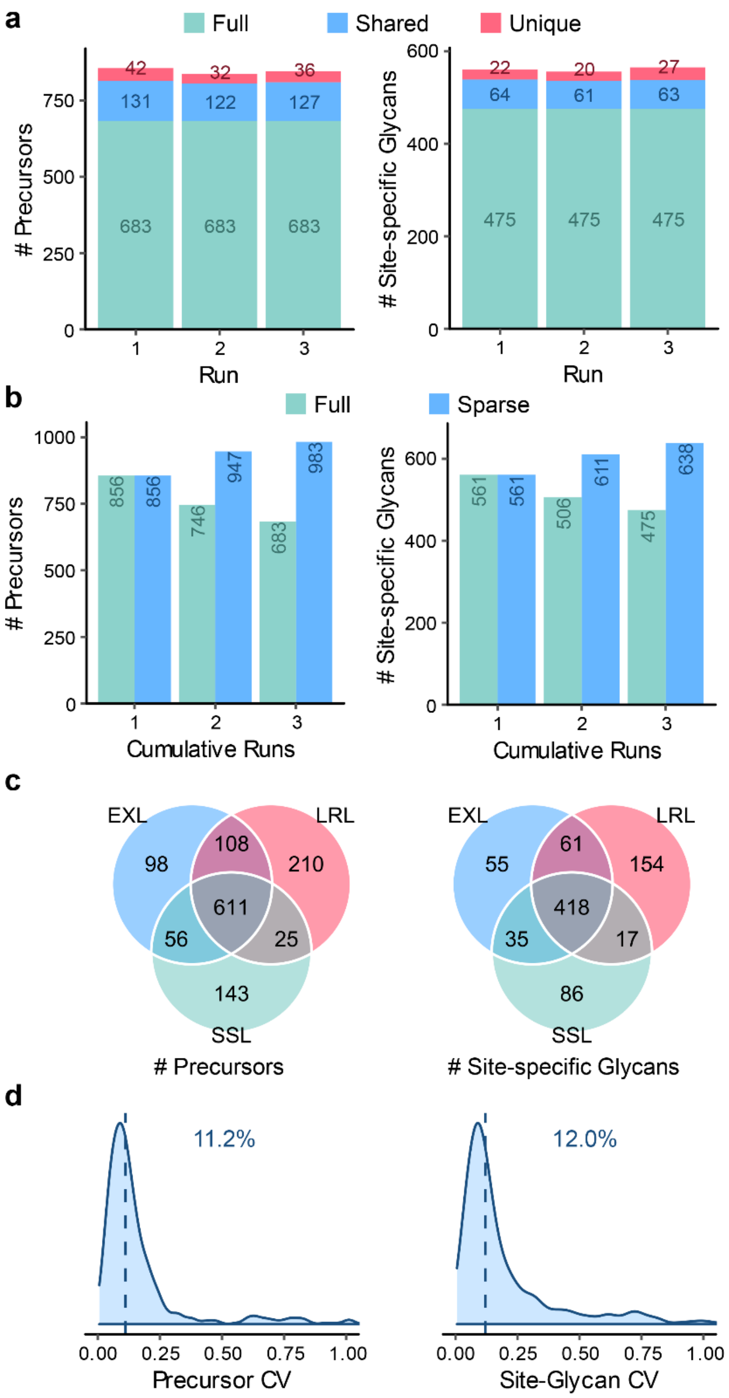


**Supplementary Fig. 23.** DIA results of the human serum sample using the extended library at the level of precursor and site-specific glycan. (**a**) Numbers of identifications per run. “Full” represents identifications observed in all the runs; “shared” represents identifications observed in 2 runs; “unique” represents identifications observed in only 1 run. (**b**) Numbers of cumulative identifications from run 1 to 3. “Full” represents identifications shared in the cumulative runs; “sparse” represents identifications observed in at least one run in the cumulative runs. (**c**) Comparison of numbers of identifications shared in >50% runs using the sample-specific library (SSL), the lab repository-scale library (LRL), and the extended library (EXL). (**d**) Coefficients of variation (CVs) of quantification results by DIA with EXL. Medians are indicated.
